# Structural basis for broad sarbecovirus neutralization by a human monoclonal antibody

**DOI:** 10.1101/2021.04.07.438818

**Authors:** M. Alejandra Tortorici, Nadine Czudnochowski, Tyler N. Starr, Roberta Marzi, Alexandra C. Walls, Fabrizia Zatta, John E. Bowen, Stefano Jaconi, Julia di iulio, Zhaoqian Wang, Anna De Marco, Samantha K. Zepeda, Dora Pinto, Zhuoming Liu, Martina Beltramello, Istvan Bartha, Michael P. Housley, Florian A. Lempp, Laura E. Rosen, Exequiel Dellota, Hannah Kaiser, Martin Montiel-Ruiz, Jiayi Zhou, Amin Addetia, Barbara Guarino, Katja Culap, Nicole Sprugasci, Christian Saliba, Eneida Vetti, Isabella Giacchetto-Sasselli, Chiara Silacci Fregni, Rana Abdelnabi, Shi-Yan Caroline Foo, Colin Havenar-Daughton, Michael A. Schmid, Fabio Benigni, Elisabetta Cameroni, Johan Neyts, Amalio Telenti, Gyorgy Snell, Herbert W. Virgin, Sean P.J. Whelan, Jesse D. Bloom, Davide Corti, David Veesler, Matteo Samuele Pizzuto

## Abstract

The recent emergence of SARS-CoV-2 variants of concern (VOC) and the recurrent spillovers of coronaviruses in the human population highlight the need for broadly neutralizing antibodies that are not affected by the ongoing antigenic drift and that can prevent or treat future zoonotic infections. Here, we describe a human monoclonal antibody (mAb), designated S2×259, recognizing a highly conserved cryptic receptor-binding domain (RBD) epitope and cross-reacting with spikes from all sarbecovirus clades. S2×259 broadly neutralizes spike-mediated entry of SARS-CoV-2 including the B.1.1.7, B.1.351, P.1 and B.1.427/B.1.429 VOC, as well as a wide spectrum of human and zoonotic sarbecoviruses through inhibition of ACE2 binding to the RBD. Furthermore, deep-mutational scanning and *in vitro* escape selection experiments demonstrate that S2×259 possesses a remarkably high barrier to the emergence of resistance mutants. We show that prophylactic administration of S2×259 protects Syrian hamsters against challenges with the prototypic SARS-CoV-2 and the B.1.351 variant, suggesting this mAb is a promising candidate for the prevention and treatment of emergent VOC and zoonotic infections. Our data unveil a key antigenic site targeted by broadly-neutralizing antibodies and will guide the design of pan-sarbecovirus vaccines.

---

Zoonotic pathogens are mainly responsible for dead-end infections but can be stably introduced in the new host sporadically as a result of adaptive mutations. Two coronaviruses (SARS-CoV and SARS-CoV-2) from distinct clades within the sarbecovirus subgenus have jumped from their natural hosts to humans in the last 20 years, with sustained person-to-person transmission^1, 2^. In contrast to SARS-CoV, which was brought under control in 2003 and disappeared in 2004 after claiming 774 lives, the emergence of SARS-CoV-2 at the end of 2019 has resulted in a global pandemic causing over 132 million infections and more than 2.8 million fatalities as of April 2021. SARS-CoV-2 genetic drift has resulted in new emerging variants of concern (VOC) characterized by higher transmissibility, immune evasion and/or disease severity such as B.1.1.7, B.1.351, B.1.429, and P.1 originally identified in the UK, South Africa, California, and Brazil, respectively^3-6^. Furthermore, recent data suggest that people infected with or vaccinated against the prototypic SARS-CoV-2 virus may have reduced protection from reinfection with these variants^7-15^. Therefore, pan-sarbecovirus countermeasures, such as vaccines and therapeutics, are needed to cope with SARS-CoV-2 evolution and to protect against future sarbecovirus emergence.

Similar to other coronaviruses, the sarbecovirus spike protein (S) mediates viral entry into host cells, represents the main target of neutralizing antibodies (nAbs) and is the focus of vaccine design. S comprises an S_1_ subunit, which recognizes host cell receptors, and an S_2_ subunit that promotes viral-cell membrane fusion^16, 17^. The S_1_ subunit includes the receptor-binding domain (RBD, also called S1^B^), which in the case of SARS-CoV and SARS-CoV-2 interacts with angiotensin-converting enzyme 2 (ACE2) to allow the virus to enter cells^1, 16, 18-20^.

A highly conserved region on the sarbecovirus RBDs, designated antigenic site II^21^, has been shown to elicit SARS-CoV and SARS-CoV-2 cross-neutralizing Abs (also defined as class 4 RBD-specific Abs)^22, 23^. However, site II is cryptic and becomes accessible only when at least two RBDs in the spike trimer adopt an open conformation and thus it is poorly immunogenic, as demonstrated by the low percentage of antibodies targeting this site in SARS-CoV-2 exposed individuals^21, 23^. Here we describe a site II-targeting mAb designated S2×259, which was isolated from an infected SARS-CoV-2-exposed subject, that possesses exceptional neutralization breadth within the sarbecovirus subgenus, including all VOC, and remarkable resistance to escape mutations. In addition, we show that S2×259 neutralizes ACE2-dependent SARS-CoV and SARS-CoV-2 related pseudoviruses with multiple mechanisms of action and that prophylactic administration of this mAb protects Syrian hamsters against SARS-CoV-2 challenge, including with the B.1.351 VOC. Overall, our findings indicate that S2×259 is a promising countermeasure to simultaneously protect against SARS-CoV-2 antigenic drift as well as emerging zoonotic sarbecoviruses and highlight the importance of RBD site II for pan-sarbecovirus vaccine design.

## Identification of a broadly neutralizing sarbecovirus mAb

To identify mAbs with potent and broad neutralizing activity against sarbecoviruses, we sorted SARS-CoV-2 S-specific (IgG) memory B cells from one COVID-19 convalescent individual 75 days after symptoms onset. We identified one mAb, designated S2×259, derived from VH1-69/D3-10/JH3 and VL1-40/JH3 genes (VH and VL 94.1% and 98.26 % identical to V germline genes, respectively) **(Table S1)**, which cross-reacted with 29 out of 30 S glycoproteins representative of all sarbecovirus clades^22, 24^ and current SARS-CoV-2 VOCs transiently expressed in ExpiCHO cells (**Fig. 1a,b**). The recognition of spikes from the divergent bat Asian-related (clade 2) and bat non-Asian (clade 3) clades highlights the exceptional breadth of this mAb (**Fig. 1a,b**).

**Fig. 1.**
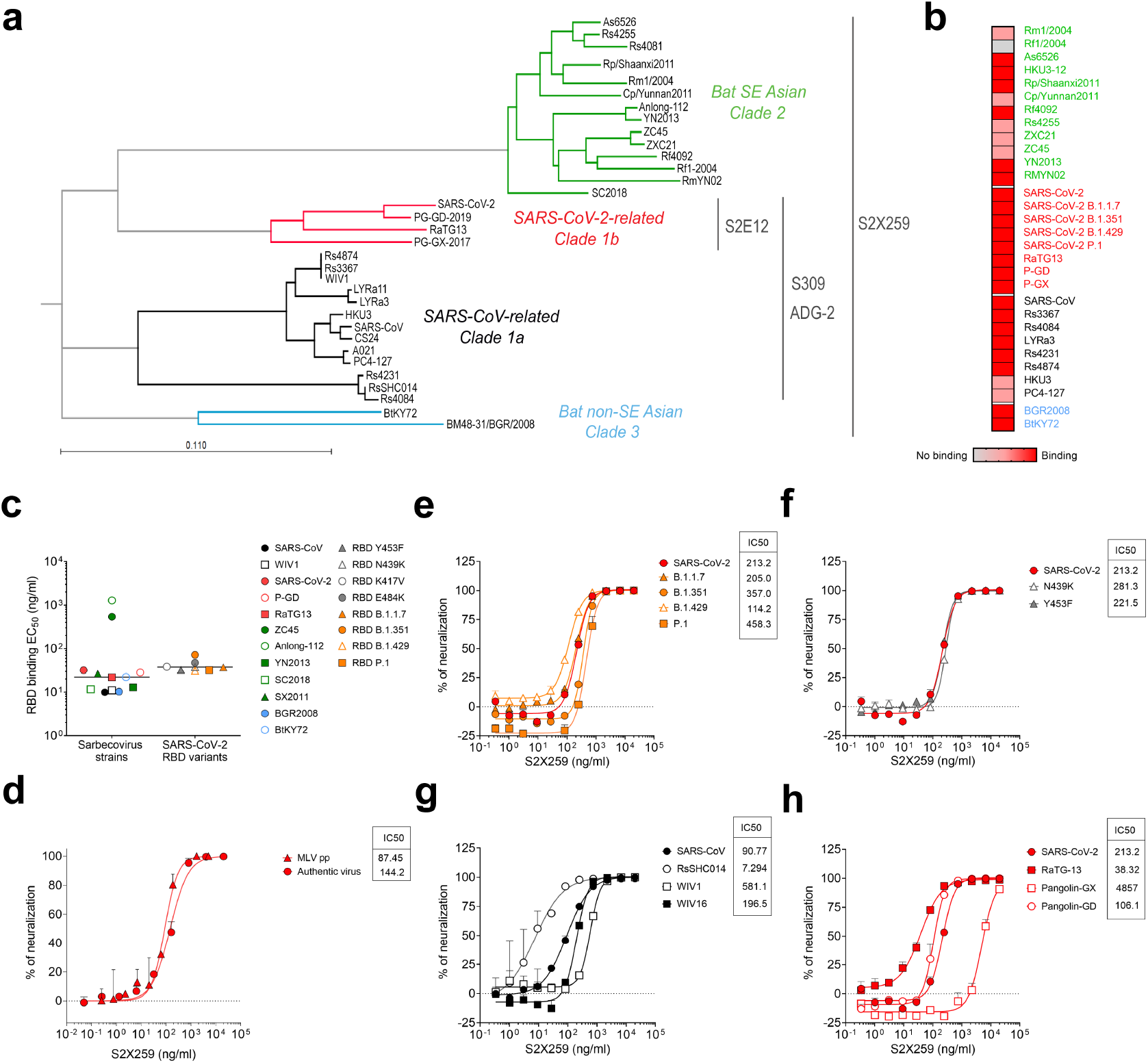
Identification of a broadly neutralizing sarbecovirus mAb. **a**, Phylogenetic tree of sarbecovirus RBDs constructed via maximum likelihood analysis of amino acid sequences retrieved from GISAID and GenBank. Cross-reactivity within the sarbecovirus subgenus for S2E12^40^, S309^30^, and ADG-2^24^ is included for comparison. **b**, Flow cytometry analysis of S2×259 cross-reactivity with a panel of 30 S glycoproteins representative of sarbecovirus clades 1a, 1b, 2, and 3 as well as SARS-CoV-2 VOCs. **c**, S2×259 binding to RBDs representative of the different sarbecovirus clades and SARS-CoV-2 variants as measured by ELISA. **d**, S2×259-mediated neutralization of SARS-CoV-2-Nluc authentic virus and SARS-CoV-2 S MLV- pseudotyped virus. **e-f** S2×259-mediated neutralization of VSV pseudotypes harbouring SARS-CoV-2 S from isolates representing the B.1.1.7, B.1.351, P.1 and B.1.429 VOC (e) as well as single RBD mutants (f). **g-h**, S2×259-mediated neutralization of VSV pseudotypes harbouring SARS-CoV-related (clade 1a, g) or SARS-CoV-2-related (clade 1b, h) S glycoproteins. One independent experiment out of at least two is shown. Error bars indicate standard deviation of duplicates or triplicates.

The cross-reactivity of S2×259 within the sarbecovirus subgenus was further confirmed using ELISA with a panel of 12 RBDs **(Fig. 1c)**. S2×259 bound tightly to all RBDs tested (EC50 < 30 ng/ml) with the exception of two bat strains ZC45 (EC50 = 539 ng/ml) and Anlong-112 (EC50 = 1281 ng/ml) (**Fig. 1c**). Surface plasmon resonance (SPR) indicated that the S2×259 Fab fragment bound to clade 1a, 1b and clade 3 RBDs with nano-to picomolar affinities, and to clade 2 RBDs with micro-to nanomolar affinities. The affinity of S2×259 for the Pangolin-Guanxi (P-GX) RBD, however, was almost 10-fold lower compared to other clade 1b RBDs (**Extended Data Fig. 1a**). The S2×259 Fab recognized both the SARS-CoV-2 prefusion-stabilized S ectodomain trimer and the isolated SARS-CoV-2 RBD with picomolar affinity albeit with a slower on-rate for binding to S presumably due to limited epitope exposure^16, 17^(**Extended Data Fig. 1a**). Finally, S2×259 binding was unaffected by several point mutants or constellation of mutations identified in circulating clinical isolates (K417V, N439K, Y453F, E484K) and in the B1.1.7 (N501Y), B.1.351 (K417N/E484K/N501Y), B.1.427/B.1.429 (L452R), and P.1 (K417T/E484K/N501Y) lineage RBDs **(Fig. 1c and Extended Data Fig. 1b)**.

We next evaluated the neutralization potency of S2×259 using authentic SARS-CoV-2-Nluc as well as murine leukemia virus (MLV) and vesicular stomatitis virus (VSV) pseudotyping systems. S2×259 potently neutralized authentic SARS-CoV-2, SARS-CoV-2 S MLV and SARS-CoV-2 S VSV with respective half-maximal inhibitory concentrations of 144 ng/mL, 87 ng/mL and 213 ng/mL **(Fig. 1d-e and Extended Data Fig. 2)**. We subsequently evaluated S2×259-mediated neutralization against VSV pseudotypes harboring SARS-CoV-2 S from the B.1.1.7, B1.351, P.1, and B.1.427/B.1.429 lineages of concern and the N439K or the Y453F mutation identified in circulating isolates^6, 25-27^. S2×259 neutralized all of these mutants without significant change in potency, in line with the identical binding affinities determined for each VOC RBD **(Fig. 1e-f and Extended Data Fig. 1b)**. S2×259 further neutralized a broad panel of ACE2-utilizing sarbecoviruses with comparable potencies, including SARS-CoV-2 Pangolin Guangdong (P-GD), RaTG13, SARS-CoV, WIV1, WIV16 and SHC014. Finally, in line with the SPR data, S2×259 neutralized P-GX S VSV pseudotype but with reduced efficiency compared to SARS-CoV-2 **(Fig. 1g-h and Extended Data Fig. 1a and 2)**.

The exceptional cross-reactivity and neutralization breadth of S2×259 against SARS-CoV-2 variants and sarbecoviruses (including viruses that have not emerged in humans) indicate this mAb is a promising candidate not only for COVID-19 prophylaxis or therapy but also for pandemic preparedness.

### Structural basis for broad sarbecovirus neutralization

To understand the molecular basis of S2×259-mediated broad neutralizing activity, we characterized a complex between the SARS-CoV-2 S ectodomain trimer and the S2×259 Fab fragment using cryo-electron microscopy (cryoEM). 3D classification of the cryoEM data showed the presence of S trimers saturated with three Fabs bound to open RBDs swung out to various extent for which we determined a consensus structure at 3.2 Å resolution **(Fig. 2a-b, Extended Data Fig. 3 and Table S2)**. We subsequently used local refinement to determine a 3.5 Å resolution map of the region corresponding to the S2×259 variable domains and RBD, which markedly improved local resolution. In parallel, we crystallized the S2×259 Fab-bound SARS-CoV-2 RBD (in the presence of the S2H97 Fab^28^) and determined a 2.65 Å resolution structure **(Fig. 2c-e and Table S3)**.

**Fig. 2.**
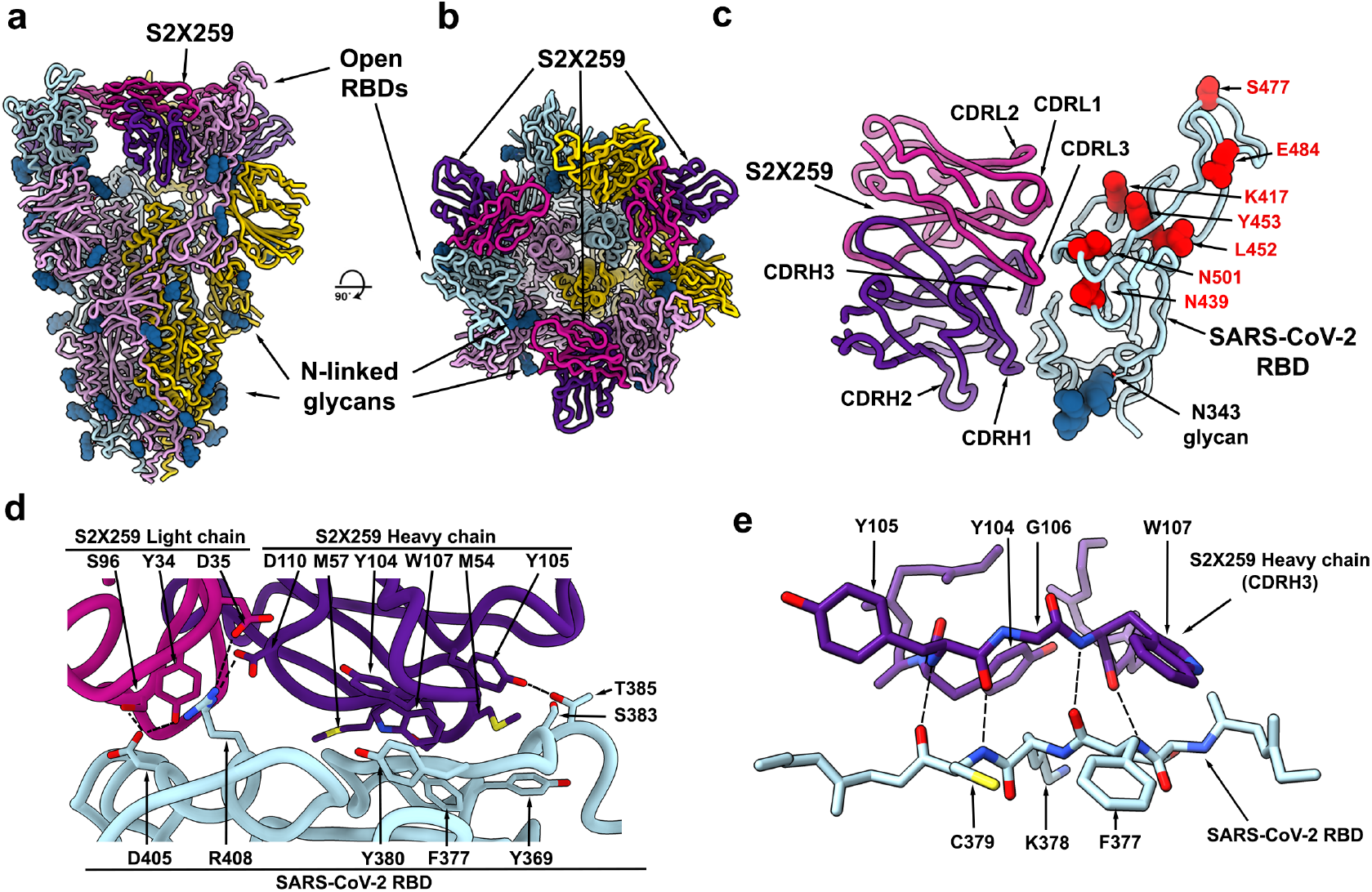
The S2×259 broadly neutralizing sarbecovirus mAb recognizes RBD antigenic site II. **a-b**, CryoEM structure of the prefusion SARS-CoV-2 S ectodomain trimer with three S2×259 Fab fragments bound to three open RBDs viewed along two orthogonal orientations. **c**. The S2×259 binding pose involving contacts with multiple RBD regions. Residues corresponding to prevalent RBD mutations are shown as red spheres. **d-e**, Close-up views showing selected interactions formed between S2×259 and the SARS-CoV-2 RBD. In panels a-e, each SARS-CoV-2 S protomer is coloured distinctly (cyan, pink and gold) whereas the S2×259 light and heavy chain variable domains are coloured magenta and purple, respectively. N-linked glycans are rendered as blue spheres in panels a-c.

S2×259 recognizes a glycan-free, cryptic epitope within antigenic site IIa which was previously defined based on the S2×35 mAb isolated from the same donor^21^. S2×259 binding requires opening of two RBDs to grant access to the Fab in the context of the S trimer **(Fig. 2a-b)**. S2×259 contacts the RBD using both heavy and light chains, which contribute approximately two thirds and one third of the ∼950 Å^2^ paratope surface buried upon binding, respectively. Specifically, S2×259 uses complementary determining regions (CDRs) H1-H3, L1 and L3 to contact RBD residues 369-386, which form two ɑ-helices and an intervening β-strand belonging to the structurally conserved RBD β-sheet, as well as the residues 404-411 and 499-508 which form a continuous surface made up of an ɑ-helix and a loop followed by an ɑ-helix, respectively.

The S2×259 epitope is conserved in circulating SARS-CoV-2 isolates and does not comprise prevalent RBD variants, such as S477N, N439K or L452R. S2×259 also circumvents residues 417 and 484, and contacts the backbone of residue N501 but not its side chain, explaining the observed high-affinity binding to the B.1.1.7, B.1.351, P.1 and B.1.429 RBDs and potent neutralization of pseudotyped viruses harbouring the S glycoprotein from these VOC **(Fig. 1c, e-f and Extended Data Fig. 1b)**. S2×259-mediated broad sarbecovirus neutralization results from the conservation of its epitope among sarbecovirus clades and from the angle of approach of the Fab which allows it to circumvent the SARS-CoV N357 glycan present in all sarbecovirus RBDs except SARS-CoV-2 (corresponding to SARS-CoV-2 residue N370) (**Extended Data Fig. 4)**. Neutralization of the SARS-CoV-2-related P-GX pseudotyped virus, however, is reduced ∼20-fold relative to SARS-CoV-2. The SARS-CoV-2 and P-GX RBDs differ at only two positions, G504N and Y508F, and the structural data indicate that the former substitution would likely dampen mAb binding through steric hindrance in agreement with the aforementioned binding and neutralization assays **(Fig. 1h and Extended Data Fig. 1a)**. The conservation of the S2×259 epitope among circulating SARS-CoV-2 variants and sarbecoviruses explain the unique cross-reactivity of this mAb across all sarbecovirus clades.

Furthermore, S2×259 binding to RBD does not affect or prevent binding by site I or site IV-targeting mAbs, also defined as class 1 and 3 RBD-specific antibodies, respectively (**Extended Data Fig. 5**), which represent the majority of antibodies currently approved or in the clinic^21, 29-31^. Therefore, S2×259 can be used in combination with these mAbs to maximize breadth against currently circulating and future variants as well as to protect against potential sarbecovirus introductions from animal reservoirs.

### S2×259 is resilient to a broad spectrum of escape mutations

Motivated by the exceptional S2×259 cross-reactivity and neutralization breadth against SARS-CoV-2 variants and genetically distant sarbecoviruses, we next evaluated if potential escape mutations could confer resistance to this mAb. Using deep-mutational scanning (DMS) with a yeast-display RBD mutant library covering all 2,034 individual amino-acid mutations that do not disrupt folding or ACE2 binding^32, 33^, we exhaustively mapped RBD mutations that escape S2×259 binding. Strikingly, S2×259 binding was strongly reduced by only a restricted number of amino acid substitutions, compared to previously described neutralizing mAbs^32^, with mutations at RBD site 504 yielding the most marked reduction in binding **(Fig. 3a and Extended Data Fig. 6a-c)**. Indeed, substitution of G504 with most other amino acids reduced S2×259 binding compared to the wildtype (Wuhan-1) RBD, highlighting the importance of this epitope residue for mAb recognition **(Fig. 3a and Extended Data Fig. 6a-c)**.

**Fig. 3.**
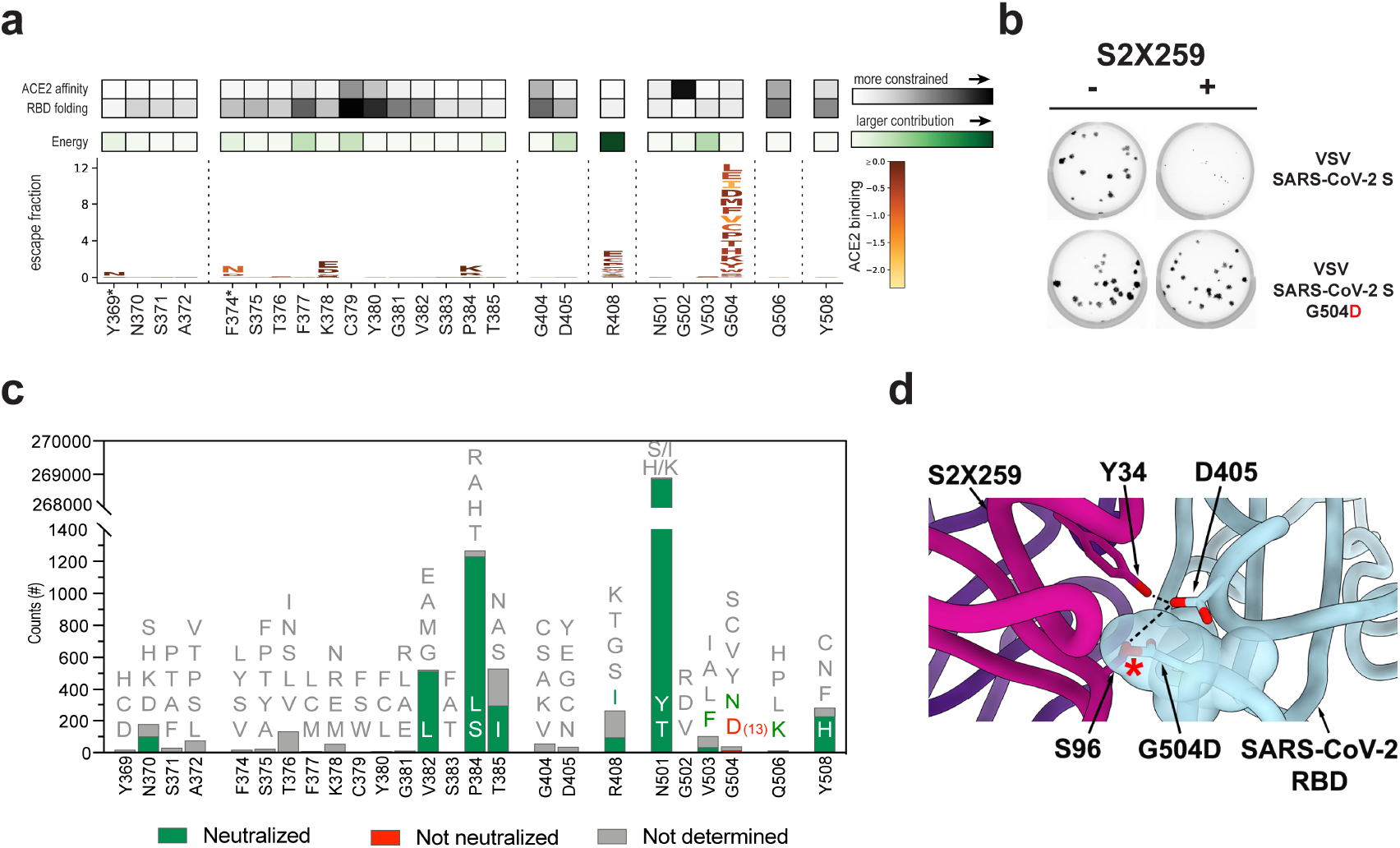
S2×259 is resilient to a broad spectrum of escape mutations. **a**, Complete mapping of mutations reducing S2×259 binding using yeast-displayed RBD deep mutational scanning (DMS). Mean mutation effect on ACE2 affinity, RBD folding, and contribution to S2×259 binding for substitutions at each position in the S2×259 epitope is reported. * mutations introducing a N-linked glycosylation sites that may not be tolerated in full spike. **b**, Plaque assays performed to validate the VSV-SARS-CoV-2 G504D mutant in Vero cells in the presence (right) or absence (left) of S2×259 in the overlay. Representative image of two independent experiments is shown. **c**, Frequency of mutants at positions encompassing S2×259 epitope based on SARS-CoV-2 genome sequences available on GISAID as of April 2021. S2×259 neutralizing activity against selected mutations is reported. **d**, Zoomed-in view of the S2×259/RBD interface showing that the G504D substitution would disrupt mAb binding due to steric hindrance.

To further explore the functional significance of the DMS data, we passaged a replication competent VSV-SARS-CoV-2 chimera (in which the native S glycoprotein was replaced by the SARS-CoV-2 S Wuhan-1 isolate gene^34, 35^) in Vero cells with S2×259 mAb in the overlay. The selective pressure imposed by S2×259 led to the deterministic emergence of viral mutants harbouring the G504D mutation, which was the sole mutation present in each of the 18 neutralization-resistant plaques sampled as a result of a single nucleotide substitution from the wildtype glycine codon (GGU) to an aspartate codon (GAU) **(Fig. 3b, Table S4 and Extended Data Fig. 6d)**. The selection of a single escape mutation suggests that the S2×259 epitope might not tolerate amino acid changes without altering viral fitness, which could be at the basis of its high conservation across the sarbecovirus subgenus.

The G504D substitution is very rare in circulating human SARS-CoV-2 isolates, as it has been detected in only 13 out of 911,641 (0.0014%) SARS-CoV-2 genome sequences available on GISAID as of April the 2^nd^ 2021 **(Fig. 3c)**. Furthermore, S2×259 efficiently neutralized VSV pseudoviruses harbouring S mutations to S2×259-contact residues found with higher frequency in clinical isolates **(Fig. 3c and Extended Data Fig. 6e)**, in agreement with the DMS results. Collectively, DMS and mAb escape selection experiments highlight the importance of position G504 for S2×259-mediated neutralization. S2×259 CDRL1 and CDRL3 contact G504 or surrounding residues, explaining that substituting glycine with any other (bulkier) residue prevents mAb binding due to steric hindrance and the potential disruption of the interactions between the Fab and residue D405, which is in close proximity to G504 (**Fig. 3d**).

These data also support the hypothesis proposed above for the markedly reduced affinity and neutralization potency of S2×259 against the P-GX RBD and pseudotyped virus which harbours the G504N mutation but maintains an aspartate at position 405 (**Fig. 1h, Extended Data Figs. 1a, 6e and 7**). However, S2×259 binding to clade 2 S bearing G504D/E mutations (e.g. Shanxi2011 or YN2013) was not abrogated (**Fig. 1c**) possibly due to substitution at the SARS-CoV-2 equivalent position 405 with a serine and to substantial deletions in the RBM (**Extended Data Fig. 7**).

Our results point to a high barrier for the emergence of S2×259 resistance mutants, which might prove essential during the next stages of the pandemic, where increasing immune pressure and continuing spread of the virus might result in the emergence of new variants.

### S2×259 blocks ACE2 engagement and protects Syrian hamsters from SARS-CoV-2 challenge

Although S2×259 targets a conserved epitope in antigenic site II, which is distinct from the RBM, some epitope residues also participate in ACE2 attachment (G502, V503 and Y505). The structural data therefore indicate that S2×259 would compete with ACE2 binding to the RBD **(Fig. 4a)**. Accordingly, we found that S2×259 blocked binding of the SARS-CoV-2 and SARS-CoV RBDs to the immobilized human recombinant ACE2 ectodomain, as measured by ELISA **(Fig. 4b)**.

**Fig. 4.**
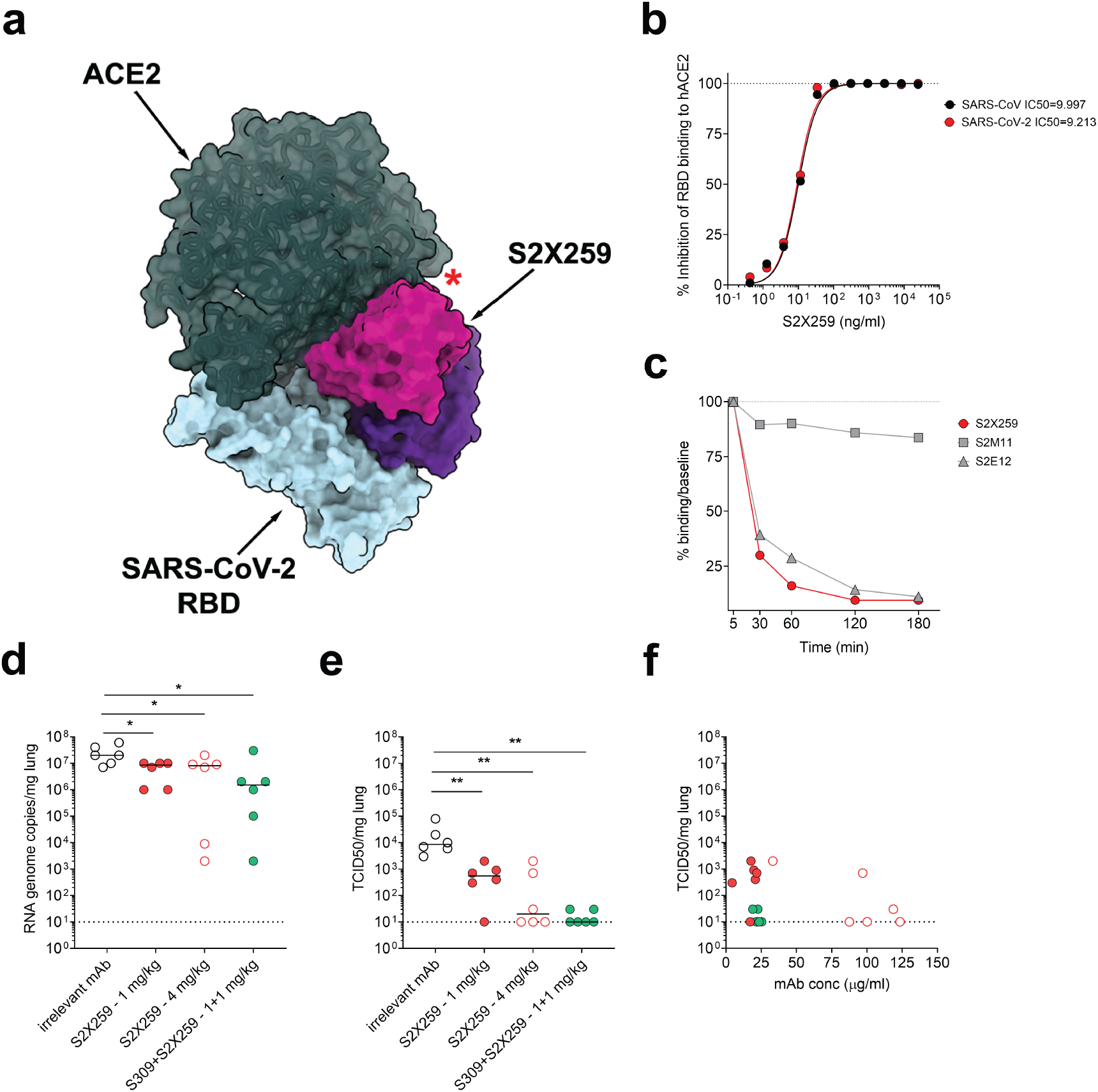
**S2×259 blocks ACE2 engagement, promotes shedding of the S_1_ subunit and protects hamsters against B.1.351 SARS-CoV-2 challenge.** **a**, S2×259 (purple/pink) and ACE2 (dark green) bind partially overlapping binding sites on the SARS-CoV-2 RBD (blue). **b**, Pre-incubation of serial dilutions of S2×259 with SARS-CoV-2 (red) or the SARS-CoV (black) RBDs prevents binding to immobilized human ACE2 (hACE2) ectodomain in ELISA. **c**, mAb-mediated S_1_ subunit shedding from cell-surface expressed SARS-CoV-2 S as determined by flow-cytometry. S2E12 was included as positive control whereas S2M11 was included as negative control. **d-e**, Quantification of viral RNA loads (d) and replicating virus titres (TCID50) (e) in the lungs of Syrian hamsters 4 days post intranasal challenge with B.1.351 SARS-CoV-2 VOC following prophylactic administration of S2×259 at 1 mg/kg (n=6), 4 mg/kg (n=6), and in combination with S309 (1+1 mg/kg, n=6). Mann–Whitney test was used for statistical analysis of significance. *p < 0.05, **p < 0.01. **f**, Correlation between concentration of mAbs measured in the serum before infection (day 0) infectious virus (TCID50) in the lung 4 days post infection. Data from one independent experiment are presented.

Since site-II targeting mAbs conformationally select for open RBDs, we assessed if S2×259 could promote shedding of the S_1_ subunit from cell-surface-expressed full-length SARS-CoV-2 S, as previously shown with RBD-specific mAbs isolated from SARS-CoV and SARS-CoV-2-exposed individuals^21, 36-39^. S2×259 binding efficiently promoted shedding of the S_1_ subunit, as was the case for the RBM-targeting S2E12 mAb but not the control mAb S2M11 which locks S in the prefusion closed state^40^ **(Fig. 4c)**. The efficient S_1_ shedding induced by S2×259 is responsible for the lack of FcγRIIa and FcγRIIIa activation, which we respectively used as proxy for Ab-dependent cellular phagocytosis (ADCP) and Ab-dependent cellular cytotoxicity (ADCC), following incubation with target cells transiently expressing full-length SARS-CoV-2 S **(Extended Data Fig. 8a-b)**. Indeed, the same tests performed using target cells transfected with a construct expressing an uncleavable pre-fusion stabilized SARS-CoV-2 S protein (unable to release the S_1_ subunit) confirmed the ability of S2×259 mAb to induce activation of FcγRIIa and FcγRIIIa in these conditions **(Extended Data Fig. 8c-d)**.

These data show that the primary mechanism of S2×259-mediated neutralization of SARS-CoV-2 and other sarbecoviruses relies on blocking viral attachment to host cell receptors through competitive inhibition of ACE2 binding. Premature triggering of S_1_ subunit shedding could inactivate viruses before encountering target cells albeit reducing activation of S2×259 mediated effector functions.

We next evaluated the prophylactic activity of S2×259 against challenge with the prototypic (Wuhan-1 related) SARS-CoV-2 in a Syrian hamster model^41^. S2×259 was administered via intraperitoneal injection 48h before intranasal challenge and the lungs of the animals were collected 4 days later for the quantification of viral RNA and replicating virus. Despite the lack of FcγRs activation *in vitro*, which has been previously ascribed with a key role for protection against SARS-CoV-2^42, 43^, S2×259 administration at 4 mg/kg reduced the amount of viral RNA detected in the lungs by 1 order of magnitude compared to hamsters receiving a control mAb **(Extended Data Fig 9a, c)**. Furthermore, greater than 2 orders of magnitude reduction in infectious virus was observed in the lungs of hamster administered with S2×259 at 4 mg/kg in comparison to the control group, **(Extended Data Fig. 9b, d)**. We next assessed the prophylactic activity of S2×259 alone and in combination with S309 against B.1.351 SARS-CoV-2 challenge^44^. S2×259 at 1 and 4 mg/kg or in combination with S309 (each at 1 mg/kg) significantly decreased the amount of viral RNA detected in the lungs at least by 1 order of magnitude in comparison to hamsters receiving a control mAb (**Fig. 4d)**. Moreover, S2×259 at 4 mg/kg and in combination with S309 (each at 1 mg/kg) completely abrogated SARS-CoV-2 B.1.351 virus replication in most of the animals **(Fig. 4e)**. The amounts of infectious viruses at day 4 detected in the group administered with only S2×259 inversely correlated with serum mAb concentration measured at the time of infection **(Fig. 4f)**. Animals receiving the mAb cocktail appeared to benefit from the contribution of S309 or from a potential synergistic activity of the two antibodies **(Fig. 4f)**.

Overall, these results show that S2×259 protects Syrian hamsters against prototypic (Wuhan-1 related) as well as B.1.351 SARS-CoV-2 replication in the lungs when prophylactically administered at 4 mg/kg. Furthermore, these data demonstrate the possibility to combine S2×259 with S309 to maximize the beneficial activity of the two mAbs at low doses against SARS-CoV-2 VOC.

## Discussion

The uncontrolled global spread of SARS-CoV-2 has led to the detection of a large number of mutations in the S glycoprotein (along with other proteins) in SARS-CoV-2 clinical isolates. Some of these variants are especially concerning such as the B.1.1.7, B.1.351 and P.1 lineages which originated in the UK, South Africa and Brazil, respectively^6, 25, 27^. The independent acquisition of identical or similar amino acid mutations among SARS-CoV-2 VOC has been shown to abrogate or reduce the neutralization potency of a large number of mAbs and vaccine-elicited sera^7-15, 45^. Moreover, the detection of a large pool of sarbecoviruses in bats and other mammals across multiple continents and the increasingly frequent interactions between humans and wildlife suggests that future cross-species transmission events are likely to occur^1, 46-49^. A critical window of opportunity after a zoonotic transmission event is the prevention of viral spread through deployment of effective countermeasures such as ring vaccination or administration of broadly neutralizing mAbs. However, approved SARS-CoV-2 therapeutics and vaccines predominantly target or elicit immunity against immunodominant but highly mutable epitopes and therefore have a limited efficacy against antigenic drift and genetically distinct zoonotic strains^45, 50^.

Strategies for eliciting broadly neutralizing and protective sarbecovirus Abs targeting the RBD have recently been proposed^22, 45^ and previous work by our group and others described mAbs now in the clinic with extended coverage within clade 1 sarbecoviruses^24, 30^. Furthermore, the recent results from the Phase III study demonstrating efficacy (i.e., 85%) of the cross-reactive VIR-7831 mAb (a derivative of the S309 antibody) paves the way for the development of additional neutralizing mAbs targeting conserved RBD epitopes that may have the dual advantage of broad coverage across sarbecoviruses and of high barrier to resistance to neutralization escape.

Here, we identified the S2×259 mAb, that targets the RBD antigenic site II and uniquely cross-reacts with all SARS-CoV-2 VOCs evaluated as well as with 30 spike trimers or RBDs from all the four sarbecovirus clades. S2×259 broadly neutralizes pseudoviruses harbouring spikes from the B.1.1.7, B.1.351., P.1 and B.1.427/B.1.429 lineages and from representative members of SARS-CoV-2 (clade 1b) and SARS-CoV (clade 1a) strains. S2×259 neutralizes WIV1 S and SHC014 S (clade 1a) pseudotyped virus that were previously suggested as potential threats to human health due to their ability to replicate in human airway cell cultures and in mice^46, 47^. Moreover, S2×259 also recognizes spikes and RBDs from bat Asian (clade 2) and non-Asian (clade 3) sarbecovirus clades. The coverage of clade 2 might be particularly important as Letko et al.^20^ showed that exogenous protease addition to pseudoviruses from this clade, which normally do not infect human cells, could result in enhanced entry in human cells. Furthermore, although clade 2 and 3 sarbecoviruses do not bind human ACE2, a recombination event following co-infection by animal and human circulating strains might result in the introduction of the ACE2-specific RBM within these viruses thus leading to ACE2-mediated entry^20, 51^. The findings that S2×259 not only neutralizes ACE2-using sarbecoviruses but also cross-reacts with sarbecoviruses that are not known to use ACE2 indicate that this mAb is a promising candidate for clinical development and could be stockpiled as part of a pandemic preparedness toolbox.

Based on the growing body of data demonstrating that Abs targeting the SARS-CoV-2 RBD account for most neutralizing activity in COVID-19 convalescent patient sera^21, 52^, we also propose that RBD-based vaccines could better promote elicitation of high titres of S2×259-like neutralizing Abs due to the enhanced accessibility of its target antigenic site compared to S-based vaccines^9, 45, 53^. We anticipate these data will guide future efforts to develop SARS-CoV-2 vaccines overcoming the emergence of variants as well as pan-sarbecovirus vaccines.

## ACKNOWLEDGEMENTS

We thank Hideki Tani (University of Toyama) for providing the reagents necessary for preparing VSV pseudotyped viruses. The authors would also like to thank also Cindy Castado and Normand Blais (GSK Vaccines) for their help in the selection of the genetically divergent sarbecoviruses used in this study. We thank Jay C. Nix for x-ray data collection, and Isaac Hoffman and Tristan I. Croll for assistance in refinement of crystal structures. This study was supported by the National Institute of General Medical Sciences (R01GM120553 to D.V.), the National Institute of Allergy and Infectious Diseases (DP1AI158186 and HHSN272201700059C to D.V.; R01AI141707 to J.D.B.), a Pew Biomedical Scholars Award (D.V.), Investigators in the Pathogenesis of Infectious Disease Awards from the Burroughs Wellcome Fund (D.V.), Fast Grants (D.V.), the Pasteur Institute (M.A.T), the Damon Runyon Cancer Research Foundation (T.N.S.) and the University of Washington Arnold and Mabel Beckman cryoEM center. J.D.B. is an Investigator of the Howard Hughes Medical Institute. Use of the Stanford Synchrotron Radiation Lightsource, SLAC National Accelerator Laboratory, is supported by the U.S. Department of Energy, Office of Science, Office of Basic Energy Sciences under Contract No. DE-AC02-76SF00515. The SSRL Structural Molecular Biology Program is supported by the DOE Office of Biological and Environmental Research, and by the National Institutes of Health, National Institute of General Medical Sciences (P30GM133894). The contents of this publication are solely the responsibility of the authors and do not necessarily represent the official views of NIGMS or NIH.

## AUTHOR CONTRIBUTIONS

Experiment Design: M.A.T., N.C., T.N.S., R.M., S.P.J.W, J.D.B., D.C., D.V., and M.S.P; Donors’

Recruitment and Sample Collection: E.C., F.B.; PBMC Sample Processing: R.M., F.Z., A.D.M., D.P., M.B.; Protein expression and purification: M.A.T, N.C., S.J., J.E.B. K.C., N.S., C.S., I.G., and E.C.; Isolation of mAbs: R.M., F.Z., A.D.M., D.P., M.B., and C.S.F.; Binding and Neutralization assays: M.A.T., A.C.W., R.M., F.Z., D.P., M.B., M.P.H., F.A.L., H.K. M.M., J.Z., and C.H.; BLI/SPR assays: A.C.W., F.Z., A.D.M, L.E.R., and E.D.; Cryo-EM Data Collection, Processing, and Model Building: M.A.T, Z.W. and D.V.; ACE2 binding inhibition and S1 shedding: D.P. and A.D.M; Evaluation of effector functions: B.G.; Deep Mutational Scanning (DMS): T.N.S., A.A., and J.D.B.; Bioinformatic analysis of virus diversity and variants: J.D.I, I.B., and A.T.; Escape mutants selection and sequencing: Z.L., and S.P.J.W.; hamster model and data analysis: R.A., S.C.F., E.V., M.A.S., F.B., J.N., D.C., and M.S.P.; Data analysis: M.A.T., N.C., T. N. S., A.C.W., R.M., J. D. B., D.C., D.V., and M.S.P.; Manuscript Writing: M.A.T., N.C., T.N.S., R.M., J.N., A.T., G.S., H.W.V, S.P.J.W, J.D.B., D.C., D.V., and M.S.P.; Supervision: F.A.L., L.E.R., C. H., M.A.S., F.B., E.C., J.N., A.T., G.S., H.W.V, S.P.J.W, J.D.B., D.C., D.V., and M.S.P.; Funding Acquisition: J. D. B., D.V.

## MATERIALS AND METHODS

### Cell lines

Cell lines used in this study were obtained from ATCC (HEK293T, Vero and Vero-E6), ThermoFisher Scientific (Expi CHO cells, FreeStyle™ 293-F cells and Expi293F™ cells) or were generated via lentiviral transduction (Expi CHO-S, HEK293T-ACE2).

### Sample donors

Samples from a SARS-CoV-2 recovered individual, designated as donor X (male, 52 years-old), was obtained 75 days after symptoms onset under study protocols approved by the local Institutional Review Boards (Canton Ticino Ethics Committee, Switzerland, the Ethical committee of Luigi Sacco Hospital, Milan, Italy). The donor provided written informed consent for the use of blood and blood components (e.g. PBMCs, sera or plasma).

### Isolation of peripheral blood mononuclear cells (PBMCs), plasma and sera

PBMCs were isolated from blood draw performed using tubes pre-filled with heparin, followed by Ficoll density gradient centrifugation. PBMCs were either used freshly along SARS-CoV2 Spike protein specific memory B cells sorting or stored in liquid nitrogen for later use. Sera were obtained from blood collected using tubes containing clot activator, followed by centrifugation and stored at −80 °C.

### B-cell isolation and recombinant S2×259 mAb production

Starting from freshly isolated PBMCs or upon cells thawing, B cells were enriched by staining with CD19 PE-Cy7 and incubation with anti-PE beads, followed by positive selection using LS columns. Enriched B cells were stained with anti-IgM, anti-IgD, anti-CD14 and anti-IgA, all PE labelled, and prefusion SARS-CoV-2 S with a biotinylated avi tag conjugated to Streptavidin Alexa-Fluor 647 (Life Technologies). SARSCoV-2 S-specific IgG+ memory B cells were sorted by flow cytometry via gating for PE negative and Alexa-Fluor 647 positive cells. Cells were cultured for the screening of positive supernatants. Antibody VH and VL sequences were obtained by RT-PCR and mAbs were expressed as recombinant human Fab fragment or as IgG1 (G1m3 allotype). ExpiCHO cells were transiently transfected with heavy and light chain expression vectors as previously described ^30^.

Affinity purification was performed on ÄKTA Xpress FPLC (Cytiva) operated by UNICORN software version 5.11 (Build 407) using HiTrap Protein A columns (Cytiva) for full length human and hamster mAbs and CaptureSelect CH1-XL MiniChrom columns (ThermoFisher Scientific) for Fab fragments, using PBS as mobile phase. Buffer exchange to the appropriate formulation buffer was performed with a HiTrap Fast desalting column (Cytiva). The final products were sterilized by filtration through 0.22 µm filters and stored at 4 °C.

### Sarbecovirus sequence analysis and SARS-CoV-2 mutant counts

Alignment and phylogenetic tree of the strains within the sarbecovirus subgenus was generated using MEGA 7.0.26 and CLC Main workbench 21.0.3 (Qiagen). The following sequences were retrieved from GISAID and NCBI: A021 (AAV97986.1); HKU3 (QND76020.1); WIV1 (AGZ48831.1); Rs3367 (AGZ48818.1); Anlong-112 (ARI44804.1); RsSHC014 (AGZ48806.1); Rs4081 (AGZ48798.1); YN2013 (AIA62330.1); Rs4874 (ATO98205.1); Rs4255 (ATO98193.1); Rs4231 (ATO98157.1); Rs4084 (ATO98132.1); ZXC21 (AVP78042.1); SC2018 (QDF43815.1); ZC45 (AVP78031.1); Rp/Shaanxi2011 (AGC74165.1); Rm1/2004 (ABD75332.1); Rf1-2004 (ABD75323.1); Rf4092 (ATO98145.1); BM48-31/BGR/2008 (YP_003858584.1); LYRa11 (AHX37558.1); RaTG13 (QHR63300.2); PC4-127 (AAU93318.1); CS24 (ABF68959.1); SARS-CoV2 (YP_009724390.1); LYRa3 (AHX37569.1); Cp/Yunnan2011 (AGC74176.1); SARS coronavirus Urbani (AAP13441.1); As6526 (ATO98108.1); BtkY72 (APO40579.1); RmYN02 (EPI_ISL_412977); Pangolin_Guangdong-2019 (EPI_ISL_410721); Pangolin-Guanxi-2017 (EPI_ISL_410539).

The viral sequences were obtained from GISAID EpiCoV project (https://www.gisaid.org/). Analysis was performed on sequences submitted to GISAID up to April 2^nd^, 2021. The spike protein sequences were either obtained directly from the protein dump provided by GISAID or, for the latest submitted sequences that were not incorporated yet in the protein dump at the day of data retrieval, from the genomic sequences with exonerate^54^ 2 2.4.0--haf93ef1_3 (https://quay.io/repository/biocontainers/exonerate?tab=tags) using protein to DNA alignment with parameters *-m protein2dna --refine full --minintron 999999 --percent 20* and using accession <u>YP_009724390.1</u> as a reference. Multiple sequence alignment of all human spike proteins was performed with mafft^55^ 7.475--h516909a_0 (https://quay.io/repository/biocontainers/mafft?tab=tags) with parameters *--auto --reorder -- keeplength --addfragments* using the same reference as above. Spike sequences that contained >10% ambiguous amino acid or that were < than 80% of the canonical protein length were discarded. A total of 923,686 sequences were used for analysis. Variants were then extracted as compared to the reference with R 4.0.2 (https://www.r-project.org/) using Biostrings 2.56.0.

### Binding to cell surface expressed Sarbecovirus S proteins by Flow Cytometry

ExpiCHO-S cells were seeded at 6 × 10^6^ cells/mL in a volume of 5 mL in a 50 mL bioreactor. Spike encoding plasmids HKU3 (QND76020.1), Rs3367 (AGZ48818.1), YN2013 (AIA62330.1), Rs4874 (ATO98205.1), Rs4255 (ATO98193.1), Rs4231 (ATO98157.1), Rs4084 (ATO98132.1), ZXC21 (AVP78042.1), ZC45 (AVP78031.1), Rp/Shaanxi2011 (AGC74165.1), Rm1/2004 (ABD75332.1), Rf1-2004 (ABD75323.1), BM48-31/BGR/2008 (YP_003858584.1), RaTG13 (QHR63300.2), PC4-127 (AAU93318.1), SARS-CoV2 (YP_009724390.1), LYRa3 (AHX37569.1), Cp/Yunnan2011 (AGC74176.1), SARS-CoV Urbani (AAP13441.1), As6526 (ATO98108.1), BtKY72 (APO40579.1), RmYN02 (EPI_ISL_412977), Pangolin_Guangdong-2019 (EPI_ISL_410721), Pangolin-Guanxi-2017 (EPI_ISL_410539), HKU3-12 (ADE34812.1) were diluted in cold OptiPRO SFM (Life Technologies, 12309-050), mixed with ExpiFectamine CHO Reagent (Life Technologies, A29130) and added to the cells. Transfected cells were then incubated at 37°C with 8% CO_2_ with an orbital shaking speed of 250 RPM (orbital diameter of 25 mm) for 42 hours. Transiently transfected ExpiCHO cells were harvested and washed two times in wash buffer (PBS 2% FBS, 2 mM EDTA). Cells were counted and distributed into round bottom 96-well plates (Corning, 3799) and incubated with 10 µg/ml of S2×259 mAb. Alexa Fluor647-labelled Goat Anti-Human IgG secondary Ab (Jackson Immunoresearch, 109-606-098) was prepared at 2 µg/ml and added onto cells after two washing steps. Cells were then washed twice and resuspended in wash buffer for data acquisition at ZE5 cytometer (Biorad).

### Recombinant protein production

Wild-type SARS-CoV-2 RBD (with N-terminal signal peptide and ‘ETGT’, and C-terminal 8xHis-tag) was expressed in Expi293F cells at 37°C and 8% CO_2_ in the presence of 10 µM kifunensine. Transfection was performed using the ExpiFectamine 293 Transfection Kit (Thermo Fisher Scientific). Cell culture supernatant was collected four days after transfection and supplemented with 10x PBS to a final concentration of 2.5x PBS (342.5 mM NaCl, 6.75 mM KCl and 29.75 mM phosphates). For crystallization, wild-type SARS-CoV-2 RBD was purified using a 5 mL HisTALON superflow cartridge (Takara Bio) followed by size exclusion chromatography using a Superdex 200 10/300 GL column pre-equilibrated in 20 mM Tris-HCl pH 7.5, 150 mM NaCl. RBD was deglycosylated with EndoH and mixed with a 1.3-fold molar excess of S2×259 Fab and S309 Fab. The complex was purified on a Superdex 200 10/300 GL column pre-equilibrated with 20 mM Tris-HCl pH 7.5, 150 mM NaCl. SARS-CoV-2 S hexapro protein, used for cryo-EM single particle studies, was expressed and purified as described before^40^.

### Enzyme-linked immunosorbent assay (ELISA)

96 half area well-plates (Corning, 3690) were coated over-night at 4°C with 25 µl of sarbecoviruses RBD proteins WIV1 (AGZ48831.1), Anlong-112 (ARI44804.1), YN2013 (AIA62330.1), SC2018 (QDF43815.1), ZC45 (AVP78031.1), Rp/Shaanxi2011 (AGC74165.1), BM48-31/BGR/2008 (YP_003858584.1), RaTG13 (QHR63300.2), SARS-CoV2 (YP_009724390.1), SARS-CoV Urbani (AAP13441.1), BtKY72 (APO40579.1), Pangolin_Guangdong-2019 (EPI_ISL_410721) and SARS-CoV-2 RBD mutants, prepared at 5 µg/ml in PBS pH 7.2. Plates were then blocked with PBS 1% BSA (Sigma-Aldrich, A3059) and subsequently incubated with mAb serial dilutions for 1 h at room temperature. After 4 washing steps with PBS 0.05% Tween 20 (PBS-T) (Sigma-Aldrich, 93773), goat anti-human IgG secondary antibody (Southern Biotech, 2040-04) was added and incubated for 1 h at room temperature. Plates were then washed 4 times with PBS-T and 4-NitroPhenyl phosphate (pNPP, Sigma-Aldrich, 71768) substrate was added. After 30 min incubation, absorbance at 405 nm was measured by a plate reader (Biotek) and data plotted using Prism GraphPad.

### MLV-based pseudotyped virus production and neutralization

To generate SARS-CoV-2 S murine leukemia virus pseudotyped virus, HEK293T cells were seeded in 10-cm dishes in DMEM supplemented with 10% FBS. The next day cells were transfected with a SARS-CoV-2 S glycoprotein-encoding plasmid harbouring the D19 C-terminal truncation ^56^, an MLV Gag-Pol packaging construct and the reporter vector pTG-Luc, using the X-tremeGENE HP DNA transfection reagent (Roche) according to the manufacturer’s instructions. Cells were then incubated at 37°C with 5% CO_2_ for 72 h. Supernatant was harvested and cleared from cellular debris by centrifugation at 400 x g, and stored at −80 °C.

For neutralization assays, Vero E6 cells were seeded into white 96-well plates (PerkinElmer) at 20,000 cells/well and cultured overnight at 37 °C with 5 % CO_2_ in 100 µl DMEM supplemented with 10% FBS and 1% penicillin/streptomycin. The next day, MLV-SARS-CoV-2 pseudovirus was activated with 10 µg/ml TPCK treated-Trypsin (Worthington Biochem) for 1 h at 37 °C. Recombinant antibodies at various concentrations were incubated with activated pseudovirus for 1 h at 37 °C. The Vero E6 cells were then washed with DMEM, and 50 µl of pseudovirus/mAbs mixes were added and incubated for 2 h at 37 °C with 5 % CO_2_. After incubation, 50 µl of DMEM containing 20% FBS and 2 % penicillin/streptomycin were added and the cells were incubated 48 h at 37 °C with 5 % CO_2_. Following these 48 h of infection, culture medium was removed from the cells and 50 µl/well of Bio-Glo (Promega) diluted 1:2 with PBS with Ca^2+^Mg^2+^ (Thermo Fisher) was added to the cells and incubated in the dark for 15 min before reading on a Synergy H1 Hybrid Multi-Mode plate reader (Biotek). Measurements were done in duplicate, RLU values were converted to percentage of neutralization and plotted with a nonlinear regression curve fit in Graph Prism.

### VSV-based pseudotype virus production and neutralization assay

SARS-CoV-2 S (CAD0240757.1), RaTG13 S (QHR63300.2), Pangolin-Guangdong S(QLR06867.1), Pangolin-Guanxi S (QIA48623.1), SARS-CoV S (YP 009825051.1), WIV1 S (AGZ48831.1), WIV16 S (ALK02457.1), RsSHCO14 S (AGZ48806.1), the VOC B.1.429 S (QTC60823) and the VOCs, N437K, Y453F, B.1.1.7 S, B.1.351 S and P.1 S with their corresponding mutations inserted in the SARS-CoV-2 S (CAD0240757.1) backbone were used to pseudotype VSV. Pseudotyped viruses were prepared using 293T cells seeded in 10-cm dishes. Briefly, cells in DMEM supplemented with 10% FBS, 1% PenStrep were transfected with the plasmid encoding for the corresponding S glycoprotein using lipofectamine 2000 (Life Technologies) following manufacturer’s indications. One day post-transfection, cells were infected with VSV (G*ΔG-luciferase) and after 2 h, infected cells were washed four times with DMEM before adding medium supplemented with anti-VSV-G antibody (I1-mouse hybridoma supernatant diluted 1 to 50, from CRL-2700, ATCC). Particles were harvested 18 h post-inoculation, clarified from cellular debris by centrifugation at 2,000 x g for 5 min and concentrated 10 times using a 30 kDa cut off membrane and used for neutralization experiments, aliquoted and frozen at −80°C until use in neutralization experiments.

For neutralization, stable 293T cells expressing ACE2^57^ in DMEM supplemented with 10% FBS, 1% PenStrep were seeded at 40,000 cells/well into clear bottom white walled 96-well plates and cultured overnight at 37°C. Twelve-point 3-fold serial dilutions of the corresponding mAb were prepared in DMEM and pseudotyped VSV viruses were added 1:1 to each mAb dilution in the presence of anti-VSV-G antibody from I1-mouse hybridoma supernatant diluted 50 times. After 45 min incubation at 37 °C, 40 µl of the mixture was added to the cells and 2 h post-infection, 40 μL DMEM was added to the cells. After 17-20 h 50 μL/well of One-Glo-EX substrate (Promega) was added to the cells and incubated in the dark for 5-10 min prior reading on a Varioskan LUX plate reader (ThermoFisher). Measurements were done in duplicate with two independent productions of pseudotyped viruses and RLU values were converted to percentage of neutralization and plotted with a nonlinear regression curve fit in Graph Prism.

### Neutralization of authentic SARS-CoV-2-Nluc virus

Neutralization of authentic SARS-CoV-2 by entry-inhibition assay was determined using SARS-CoV-2-Nluc, an infectious clone of SARSCoV-2 (based on strain 2019-nCoV/USA_WA1/2020) which encodes nanoluciferase in place of the viral ORF7 and demonstrated comparable growth kinetics to wildtype virus ^58^. Vero E6 cells were seeded into black-walled, clear-bottom 96-well plates at 2 x 104 cells/well and cultured overnight at 37 °C. The next day, 9-point 4-fold serial dilutions of mAbs were prepared in infection media (DMEM + 10% FBS). SARS-CoV-2-Nluc was diluted in infection media at a final MOI of 0.01 PFU/cell, added to the mAb dilutions and incubated for 30 minutes at 37 °C. Media was removed from the Vero E6 cells, mAb-virus complexes were added and incubated at 37 °C for 24 hours. Media was removed from the cells, Nano-Glo luciferase substrate (Promega) was added according to the manufacturer’s recommendations, incubated for 10 minutes at room temperature and the luciferase signal was quantified on a VICTOR Nivo plate reader (Perkin Elmer).

### Affinity determination by Surface Plasmon Resonance (SPR)

SPR binding measurements were performed using a Biacore T200 instrument using either anti-AviTag pAb (for capturing S proteins) or StrepTactin XT (for capturing RBDs) covalently immobilized on CM5 chips. Running buffer was Cytiva HBS-EP+ (pH 7.4). All measurements were performed at 25 °C. S2×259 Fab concentrations were 11, 33, 100, and 300 nM run as single-cycle kinetics. Double reference-subtracted data were fit to a 1:1 binding model using Biacore Evaluation software, which yields an “apparent KD” for the S-binding data because the kinetics also reflect S conformational dynamics. For SARS-CoV-2 S the dissociation rate was too slow to fit, so the K_D_,app is reported as an upper limit. The K_D_ above 1 µM is approximate and was determined from a fit where Rmax was set to a constant based on theoretical Rmax.

### Affinity determination using bio-layer interferometry

Biotinylated RBD (WT, N501Y, K417N-E484K-N501Y, or K417T-E484K-N501Y) were immobilized at 5 ng/µL in undiluted 10X Kinetics Buffer (Pall) to SA sensors until a load level of 1.1nm. A 1:3 dilution series of Fab in undiluted kinetics buffer starting at 10nM was used for 600 seconds association prior to 600 second dissociation to determine protein-protein affinity. The data were baseline subtracted and the plots fitted using the Pall FortéBio/Sartorius analysis software (version 12.0). Data were plotted in Prism.

### Competition assay by Biolayer Interferometry (BLI)

BLI was used to assess S2×259 competition with S309 and S2E12 using an Octet Red96 (ForteBio). All reagents were prepared in kinetics buffer (PBS 0.01% BSA) at the indicated concentrations. His-tagged SARS-CoV-2 RBD was prepared at 8 µg/ml and loaded on pre-hydrated anti-penta-HIS biosensors (Sartorius) for 2.5 min. Biosensors were then moved into a solution containing S2×259 mAb and association recorded for 7 min. A second association step was subsequently performed into S2×259 (as control), S309 and S2E12 mAbs solutions at 20 µg/ml and recorded for 7 min. Response values were exported and plotted using GraphPad Prism.

### Cell-surface mAb-mediated S_1_ shedding

CHO cells stably expressing wild-type SARS-CoV-2 S were resuspended in wash buffer (PBS 1 % BSA, 2 mM EDTA) and treated with 10 µg/mL TPCK-trypsin (Worthington Biochem) for 30 min at 37°C. Cells were then washed and distributed into round bottom 96-well plates (90,000 cells/well). S2×259 was added to cells at 15 µg/mL final concentration for 180 min at 37 °C. Cells were collected at different time points (5, 30, 60, 120 and 180), washed with wash buffer at 4 °C, and incubated with 1.5 µg/mL secondary goat anti-human IgG, Fc fragment specific (Jackson ImmunoResearch) on ice for 20 min. Cells were washed and resuspended in wash buffer and analyzed with ZE5 FACS (Bio-rad).

### Measurement of Fc-effector functions

S2×259-dependent activation of human FcγRIIIa was performed with a bioluminescent reporter assay. ExpiCHO cells transiently expressing full-length wild-type SARS-CoV-2 S (target cells) or full-length prefusion stabilized SARS-CoV-2 S, which harbours the 2P mutation and S1/S2 furin cleavage site mutation (RRARS to SGAG) as previously described^16^, were incubated with different amounts of mAbs. After a 15-minute incubation, Jurkat cells stably expressing FcγRIIIa receptor (V158 variant) or FcγRIIa receptor (H131 variant) and NFAT-driven luciferase gene (effector cells) were added at an effector to target ratio of 6:1 for FcγRIIIa and 5:1 for FcγRIIa. Signaling was quantified by the luciferase signal produced as a result of NFAT pathway activation. Luminescence was measured after 20 hours of incubation at 37°C with 5% CO2 with a luminometer using the Bio-Glo-TM Luciferase Assay Reagent according to the manufacturer’s instructions (Promega).

### In vivo mAb testing using a Syrian hamster model

KU LEUVEN R&D has developed and validated a SARS-CoV-2 Syrian Golden hamster infection model ^41, 44^. The SARS-CoV-2 Wuhan (BetaCov/Belgium/GHB-03021/2020-EPI ISL 109 407976|2020-02-03) and B.1.351 (hCoV105 19/Belgium/rega-1920/2021; EPI_ISL_896474, 2021-01-11) isolates used in this study, were recovered from nasopharyngeal swabs taken from a RT-qPCR confirmed asymptomatic patient who returned from Wuhan, China in February 2020 and from a patient with respiratory symptoms returned to Belgium in January 2021, respectively. A close relatedness with the prototypic Wuhan-Hu-1 2019 SARS-CoV-2 and with B.1.351 lineage was confirmed by sequencing and phylogenetic analysis. Infectious viruses were isolated by serial passaging on Vero E6 cells and passage 6 for SARS-CoV-2 Wuhan and passage 2 for B.1.351 viruses were used for the study. The titre of the virus stock was determined by end-point dilution on Vero E6 cells by the Reed and Muench method ^59^. This work was conducted in the high-containment A3 and BSL3+ facilities of the KU Leuven Rega Institute (3CAPS) under licenses AMV 30112018 SBB 219 2018 0892 and AMV 23102017 SBB 219 20170589 according to institutional guidelines.

Syrian hamsters (Mesocricetus auratus) were purchased from Janvier Laboratories and were housed per two in ventilated isolator cages (IsoCage N Biocontainment System, Tecniplast) with ad libitum access to food and water and cage enrichment (wood block). Housing conditions and experimental procedures were approved by the ethical committee of animal experimentation of KU Leuven (license P065-2020). 6-10 week-old female hamsters were administered by intraperitoneal injection with S2×259 mAb at 1 mg/kg and 4 mg/kg 48 hours before intranasal infection with 1.89×106 TCID50 in 50 µl inoculum. Hamsters were monitored for appearance, behavior and weight. At day 4 post infection hamsters were euthanized by intraperitoneal injection of 500 μL Dolethal (200 mg/mL sodium pentobarbital, Vétoquinol SA). Lungs were collected, homogenized using bead disruption (Precellys) in 350 µL RLT buffer (RNeasy Mini kit, Qiagen) and centrifuged (10,000 rpm, 5 minutes, 4°C) to pellet the cell debris. RNA was extracted using a NucleoSpin kit (Macherey-Nagel) according to the manufacturer’s instructions. RT-qPCR was performed on a LightCycler96 platform (Roche) using the iTaq Universal Probes One-Step RTqPCR kit (BioRad) with N2 primers and probes targeting the nucleocapsid ^41^. Standards of SARS-CoV-2 cDNA (IDT) were used to express viral genome copies per mg tissue or per mL serum. To quantify infectious SARS-CoV-2 particles, endpoint titrations were performed on confluent Vero E6 cells in 96-well plates. Viral titres were calculated by the Reed and Muench method ^59^ and were expressed as 50% tissue culture infectious dose (TCID50) per mg tissue.

### Evaluation of escape mutants via deep mutational scanning (DMS)

A previously described deep mutational scanning approach^32^ was used to identify RBD mutations that escape S2×259 binding. Briefly, duplicate libraries containing virtually all possible amino acid changes compatible with ACE2 binding and RBD folding within the Wuhan-Hu-1 SARS-CoV-2 RBD sequence were expressed on the surface of yeast^32, 33^. Libraries were labelled at 59 ng/mL S2×259 antibody (the EC90 for binding to yeast-displayed SARS-CoV-2 RBD determined in isogenic pilot binding experiments), and fluorescence-activated cell sorting (FACS) was used to select RBD+ cells that exhibit reduced antibody binding as previously described^32, 60^. See FACS plots in Extended Data Fig. 6a. Libraries were sequenced before and after selection to determine per-mutation escape fractions as previously described^60^. Experiments were performed in duplicate with independently generated mutant libraries (correlations shown in Extended Data Fig. 6b), and we report the average mutant escape fraction across the duplicates. Raw escape fractions are available on GitHub: https://github.com/jbloomlab/SARS-CoV-2-RBD_MAP_Vir_mAbs/blob/main/results/supp_data/s2×259_raw_data.csv. Detailed steps of analysis and code for the deep mutational scanning selections are available on GitHub: https://github.com/jbloomlab/SARS-CoV-2-RBD_MAP_Vir_mAbs.

### Selection of SARS-CoV-2 monoclonal antibody escape mutants (MARMS)

VSV-SARS-CoV-2 chimera was used to select for SARS-CoV-2 S monoclonal antibody resistant mutants (MARMS) previously described (^34, 35^). Briefly, MARMS were recovered by plaque isolation on Vero cells with the indicated mAb in the overlay. The concentration of mAb in the overlay was determined by neutralization assays at a multiplicity of infection (MOI) of 100. Escape clones were plaque-purified on Vero cells in the presence of mAb, and plaques in agarose plugs were amplified on MA104 cells with the mAb present in the medium. Viral stocks were amplified on MA104 cells at an MOI of 0.01 in Medium 199 containing 2% FBS and 20 mM HEPES pH 7.7 (Millipore Sigma) at 34°C. Viral supernatants were harvested upon extensive cytopathic effect and clarified of cell debris by centrifugation at 1,000 x g for 5 min. Aliquots were maintained at - 80°C. Viral RNA was extracted from VSV-SARS-CoV-2 mutant viruses using RNeasy Mini kit (Qiagen), and S was amplified using OneStep RT-PCR Kit (Qiagen). The mutations were identified by Sanger sequencing (GENEWIZ). Their resistance was verified by subsequent virus infection in the presence or absence of antibody. Briefly, Vero cells were seeded into 12 well plates for overnight. The virus was serially diluted using DMEM and cells were infected at 37°C for 1 h. Cells were cultured with an agarose overlay in the presence or absence of mAb at 34°C for 2 days. Plates were scanned on a biomolecular imager and expression of eGFP is show at 48 hours post-infection.

### Crystallization, data collection, structure determination and analysis

Crystals of the SARS-CoV-2-RBD-S2×259-S2H97 Fab complex were obtained at 20°C by sitting drop vapor diffusion. A total of 200 nL complex at 5.7 mg/ml were mixed with 200 nL mother liquor solution containing 0.12 M Monosaccharides mix, 20% (v/v) Ethylene glycol, 10% (w/v) PEG 8000, 0.1 M Tris (base)/bicine pH 8.5, 0.02 M Sodium chloride, 0.01 M MES pH 6 and 3% (v/v) Jeffamine ED-2003. Crystals were flash frozen in liquid nitrogen. Data were collected at Beamline 9-2 of the Stanford Synchrotron Radiation Lightsource facility in Stanford, CA. Data were processed with the XDS software package^61^ for a final dataset of 2.65 Å in space group P21. The RBD-S2×259-S2H97 Fab complex structure was solved by molecular replacement using phaser^62^ from a starting model consisting of SARS-CoV-2 RBD (PDB ID: 7JX3) and homology models of the S2×259 and S2H97 Fabs built using the Molecular Operating Environment (MOE) software package from the Chemical Computing Group (https://www.chemcomp.com). Several subsequent rounds of model building and refinement were performed using Coot^63^, ISOLDE^64^, Refmac5^65^, and MOE (https://www.chemcomp.com), to arrive at a final model for the ternary complex.

### CryoEM sample preparation, data collection and data processing

Recombinantly expressed and purified fab S2×259 and SARS-CoV-2 S hexapro were incubated at 1 mg/ml with a 1.2 molar excess of fab at 4**°**C during 1 hr. Three microliters of the complex mixture were loaded onto freshly glow discharged R 2/2 UltrAuFoil grids (200 mesh) or lacey grids covered with a thin layer of manually added carbon, prior to plunge freezing using a vitrobot MarkIV (ThermoFisher Scientific) with a blot force of 0 and 7-7.5 sec blot time (for the UltrAuFoil grids) or with a blot force of −1 and 2.5 sec blot time (for the lacey thin carbon grids) at 100 % humidity and 21**°**C.

Data were acquired on a FEI Titan Krios transmission electron microscope operated at 300 kV and equipped with a Gatan K2 Summit direct detector and Gatan Quantum GIF energy filter, operated in zero-loss mode with a slit width of 20 eV. Automated data collection was carried out using Leginon^66^ at a nominal magnification of 130,000x with a super-resolution pixel size of 0.525 Å. The dose rate was adjusted to 8 counts/pixel/s, and each movie was fractionated in 50 frames of 200 msec. Two datasets were collected from UltrAuFoil grids with the stage tilted 30° and 55° to circumvent particle preferential orientation. The third dataset was collected on lacey grids covered with a thin layer of carbon. The three datasets, with a total of 6786 micrographs, were collected with a defocus range comprised between −0.8 and −2 μm. For each dataset, movie frame alignment, estimation of the microscope contrast-transfer function parameters, particle picking and extraction were carried out using Warp^67^. Particle images were extracted with a box size of 800 pixels^2^ and binned to 400 yielding a pixel size of 1.05 Å. The three datasets were merged and two rounds of reference-free 2D classification were performed using cryoSPARC^68^. Subsequently, one round of 3D classification with 50 iterations, using PDB 6VXX as initial model, was carried out using Relion^69, 70^ without imposing symmetry. 3D refinements were carried out using non-uniform refinement^71^. Particle images from each dataset were subjected to Bayesian polishing^72^ using Relion before merging them to perform another round of non-uniform refinement in cryoSPARC followed by per-particle defocus refinement and again non-uniform refinement. To improve the density of the S/S2×259 interface, the particles were symmetry-expanded and subjected to a Relion focus 3D classification without refining angles and shifts using a soft mask encompassing the RBD and S2×259 variable domains. Particles belonging to classes with the best resolved local density were selected and subjected to local refinement using cryoSPARC. Local resolution estimation, filtering, and sharpening were carried out using CryoSPARC. Reported resolutions are based on the gold-standard Fourier shell correlation (FSC) of 0.143 criterion and Fourier shell correlation curves were corrected for the effects of soft masking by high-resolution noise substitution^73^.

### CryoEM model building and analysis

UCSF Chimera^74^ and Coot^63^ were used to fit atomic models (PDB 6VXX or PDB 6VYB) into the cryoEM maps and the Fab variable domains were manually built. S2E12 was built in the locally refined map and subsequently validated using the Fab crystal structure. Models were refined and relaxed using Rosetta using both sharpened and unsharpened maps^75, 76^. Validation used Phenix^77^, Molprobity^78^ and Privateer^79^. Figures were generated using UCSF ChimeraX^80^.

**Extended Data Fig. 1.**
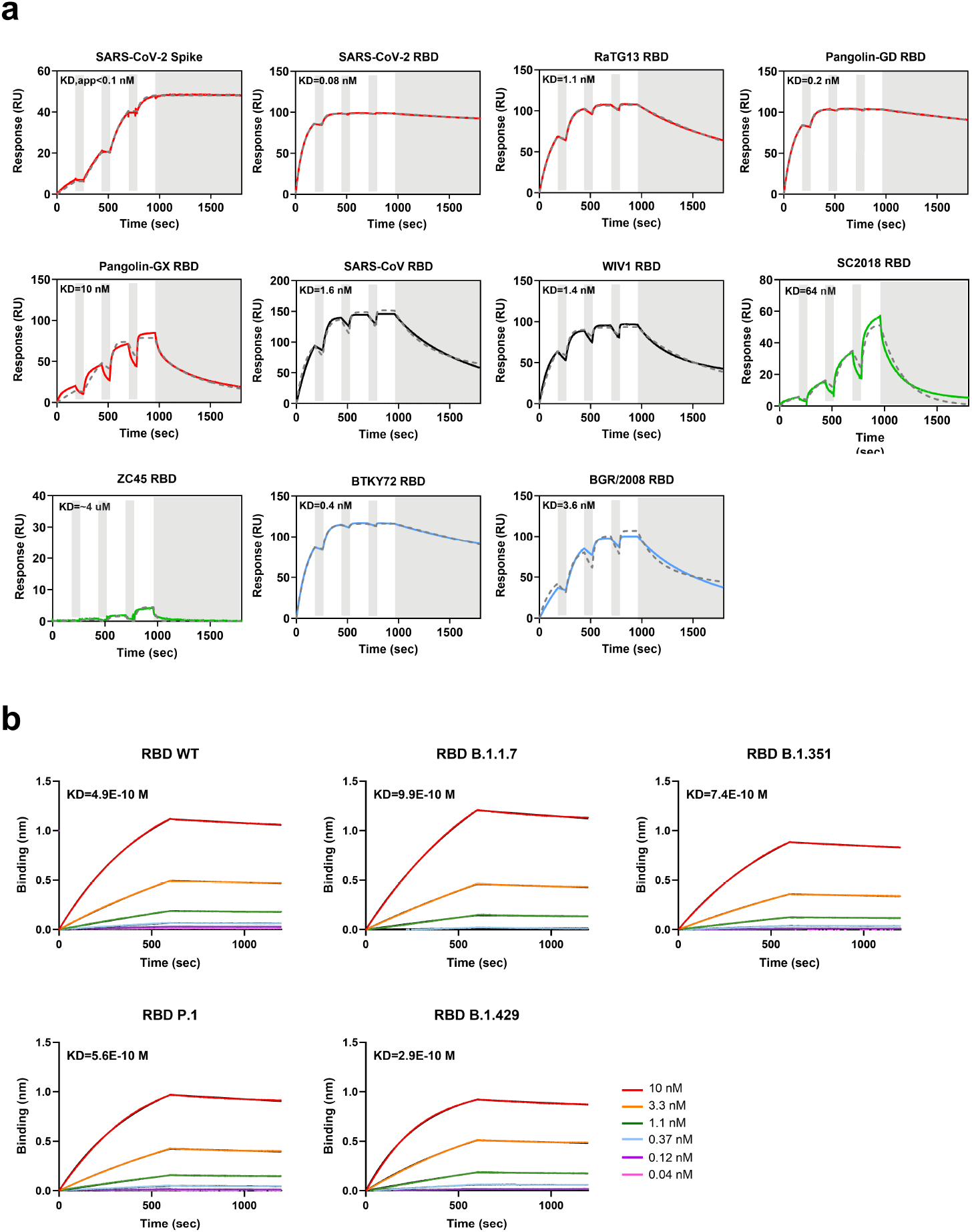
**S2×259 Fab binding to recombinant sarbecovirus RBDs, prefusion SARS-CoV-2 S ectodomain trimer and RBD variants.** **a**, S or RBD antigens were captured on the sensor chip surface and binding to S2×259 Fab at 11, 33, 100, and 300 nM was monitored successively, in single-cycle kinetics format, by surface plasmon resonance. All data have been fit to a 1:1 binding model and the equilibrium dissociation constant (K_D_) is reported. For the S-binding data, we report an apparent K_D_ (K_D,app_) since kinetics are affected by conformational dynamics between open and closed RBD states. The colouring scheme matches the phylogenetic tree in Figure 1a. **b**, Biolayer interferometry binding analysis of the S2×259 Fab to wildtype or VOC SARS-CoV-2 biotinylated RBDs immobilized at the surface of SA biosensors.

**Extended Data Fig. 2.**
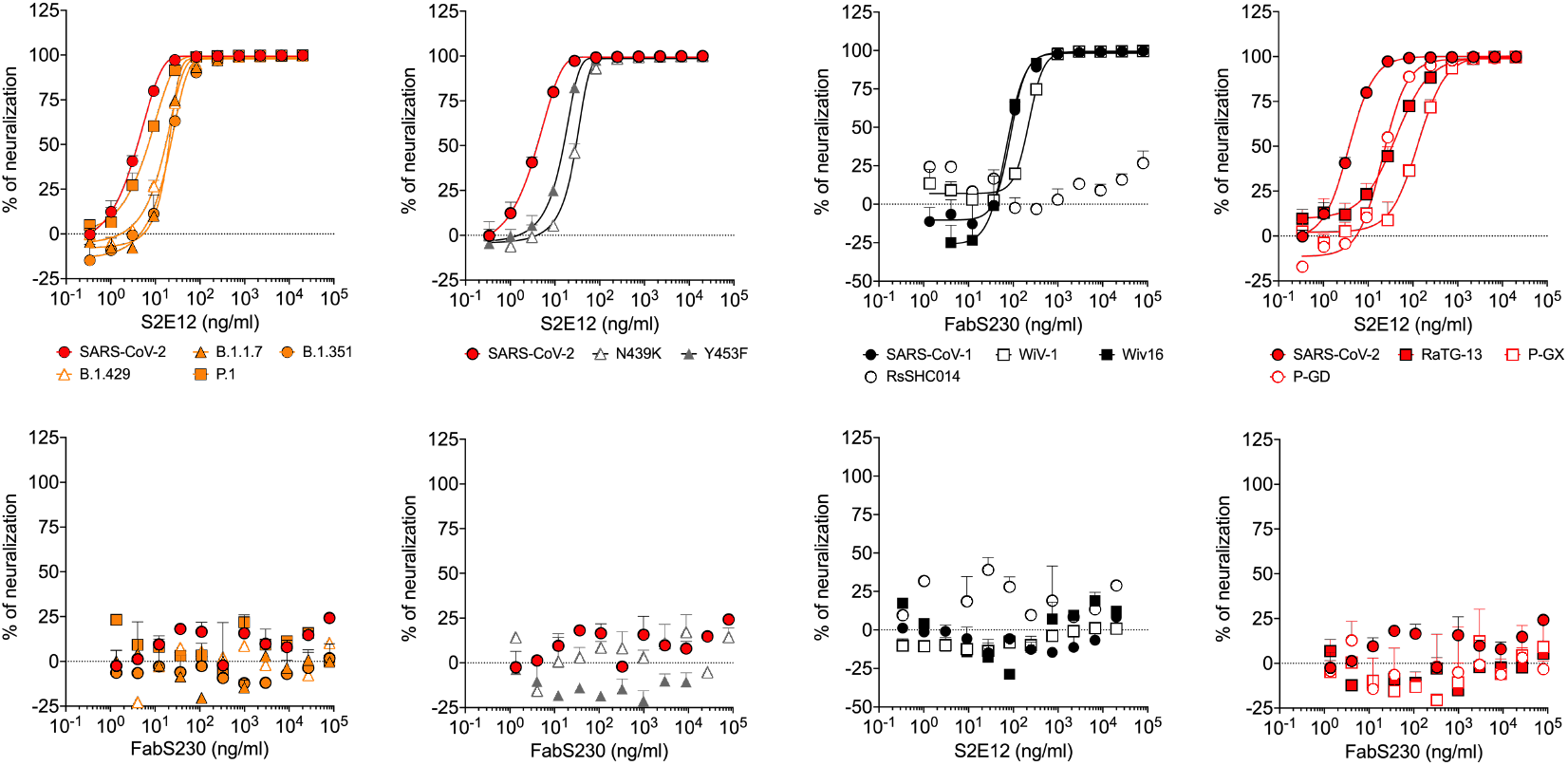
Neutralization tests with control mAbs. S2E12 and S230 neutralizing activity against VSV pseudotypes harbouring the S glycoprotein of SARS-CoV- and SARS-CoV-2 related strains and VOC.

**Extended Data Fig. 3.**
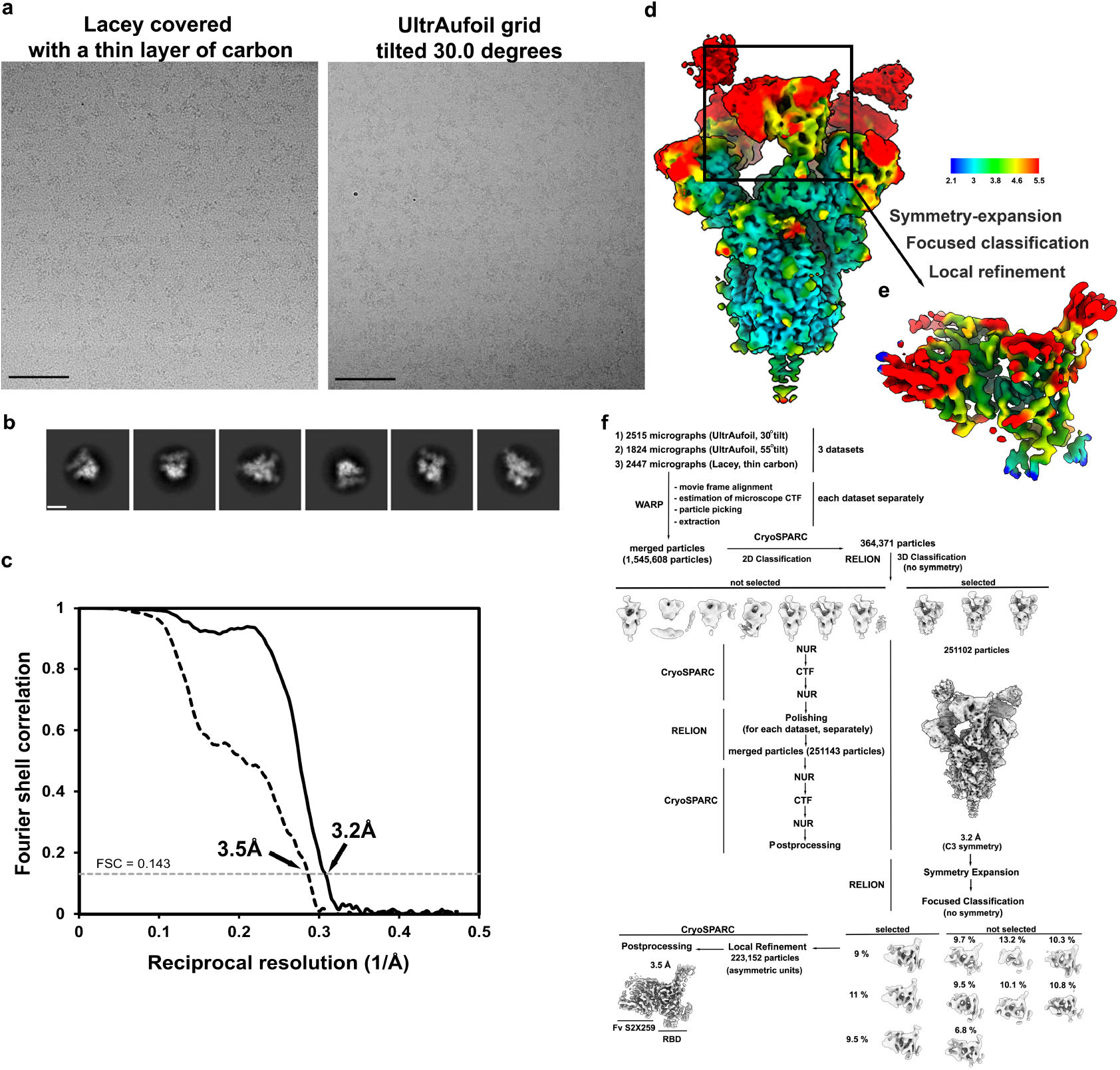
CryoEM data processing and validation of S2×259-bound SARS-CoV-2 S datasets. **a-b**, Representative electron micrographs (a) and class averages (b) of SARS-CoV-2 S in complex with the S2×259 Fab. Scale bar of the micrograph: 500 Å. Scale bar of the class averages: 100 Å. **c**, Gold-standard Fourier shell correlation curves for the S trimer bound to three S2×259 Fabs (solid black line) and the locally refined reconstruction of the RBD/S2×259 variable domains (dashed black line). The 0.143 cut-off is indicated by a horizontal dashed grey line. **d-e**, Local resolution map for the open S trimer bound to three S2×259 (d) and the locally refined reconstruction of the RBD/S2×259 variable domains (e). **f**, CryoEM data processing flow-chart.

**Extended Data Fig. 4.**
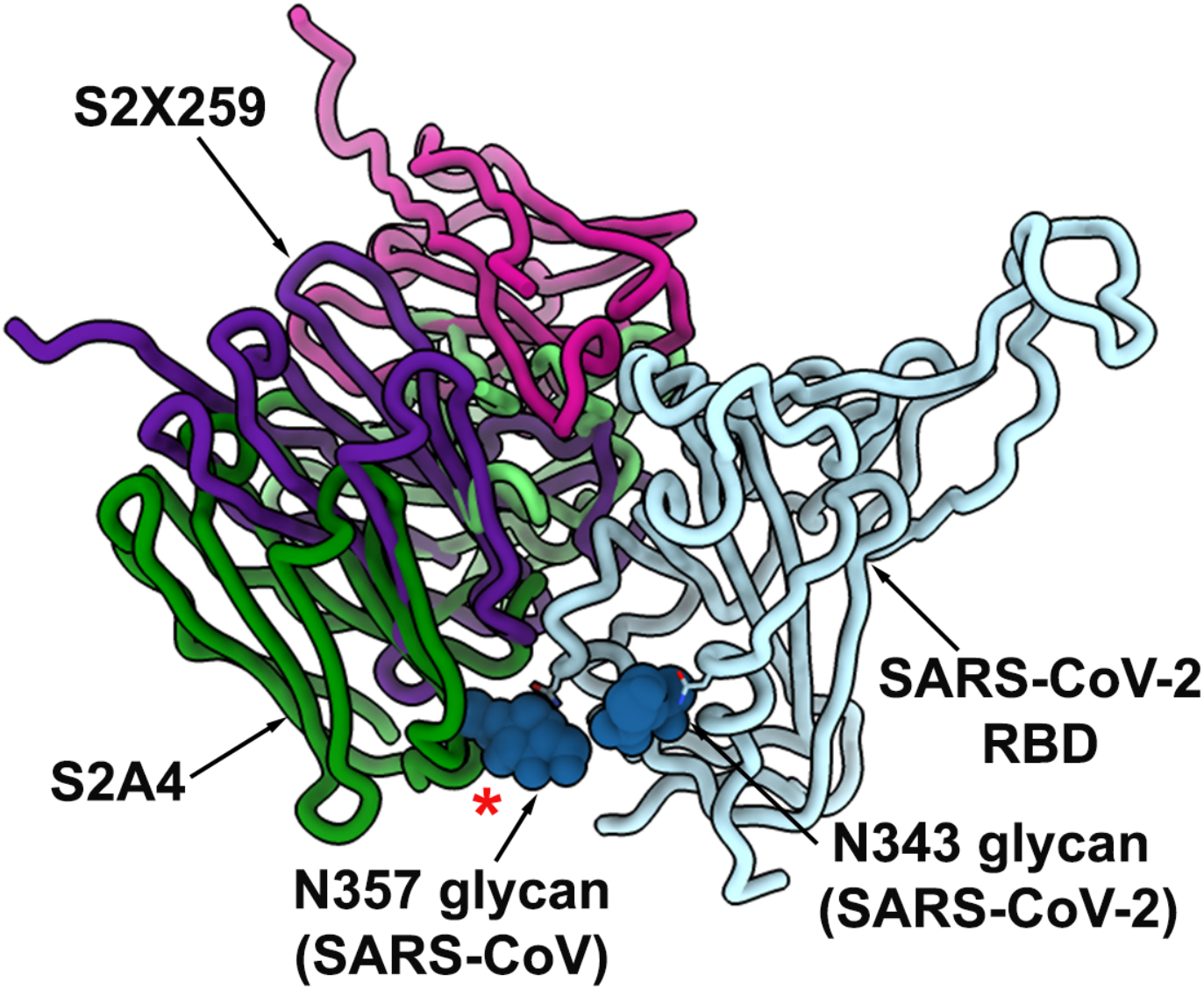
**The S2×259 angle of approach for binding to the SARS-CoV-2 RBD allows to circumvent the SARS-CoV N357 glycan present in all sarbecovirus RBDs except SARS-CoV-2**. Ribbon diagram showing a superimposition of the S2×259-bound and S2A4-bound (PDB 7JVA) SARS-CoV-2 RBD^21^. The SARS-CoV glycan at position N357 was modelled based on the S230-bound SARS-CoV S structure (PDB 6NB6^37^) and is predicted to sterically hinder S2A4 binding (red star) but not S2×259. The mAb light and heavy chains are coloured magenta and purple (S2×259) or light and dark green (S2A4). N-linked glycans are rendered as blue spheres.

**Extended Data Fig. 5.**
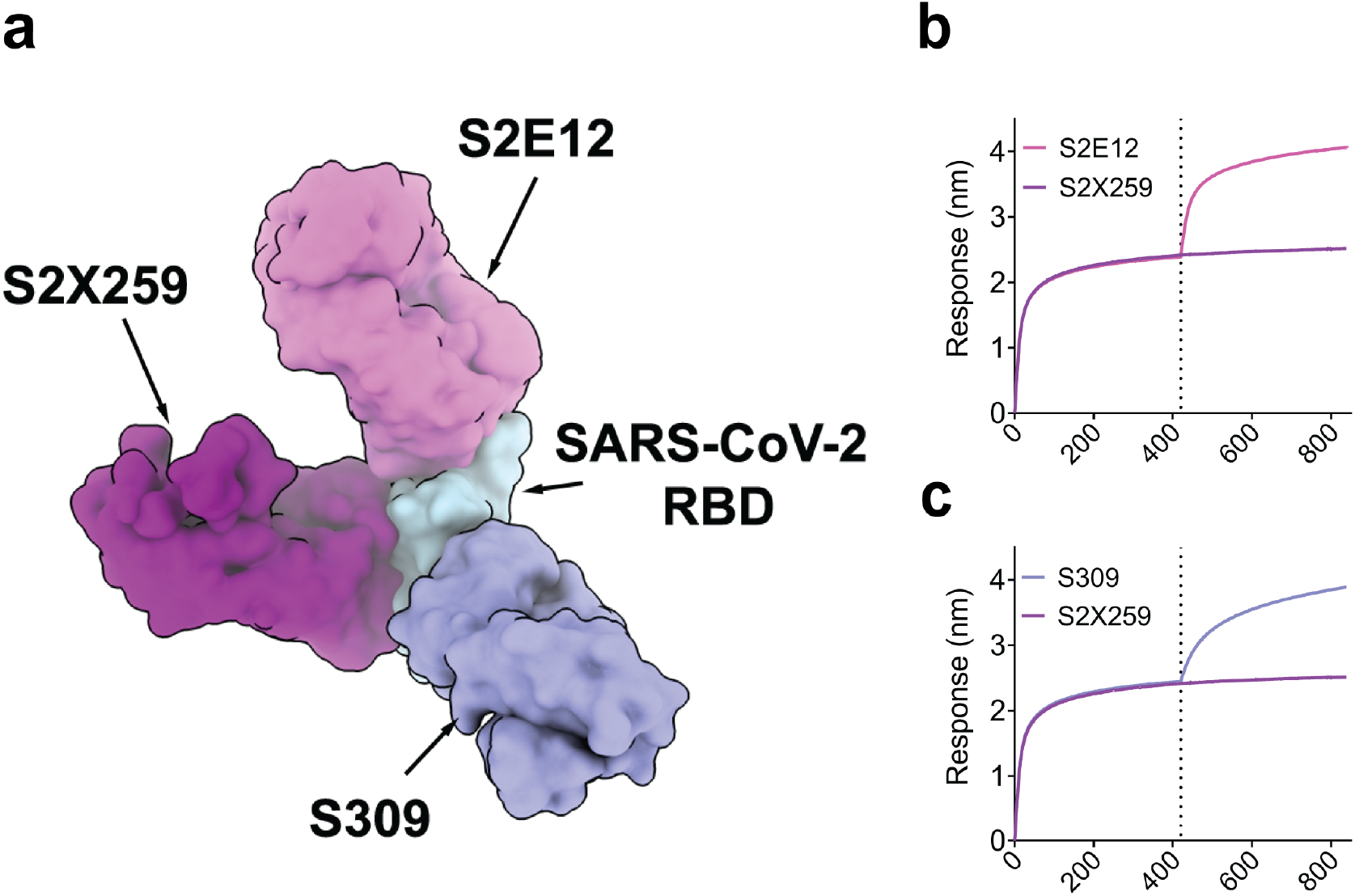
S2×259 allows combination with site I and IV-targeting mAbs. **a**, View of site I-targeting S2E12^40^ (pink), site II-targeting S2×259 (magenta), and site IV-targeting S309^30^ (purple) mAb binding to SARS-CoV-2 RBD (light blue). **b-c**, Competition binding assays for S2×259 vs site I-targeting S2E12 (b) and site IV-targeting S309 (c) mAbs on SARS-CoV-2 RBD as measured by biolayer interferometry. One independent experiment out of two is shown.

**Extended Data Fig. 6.**
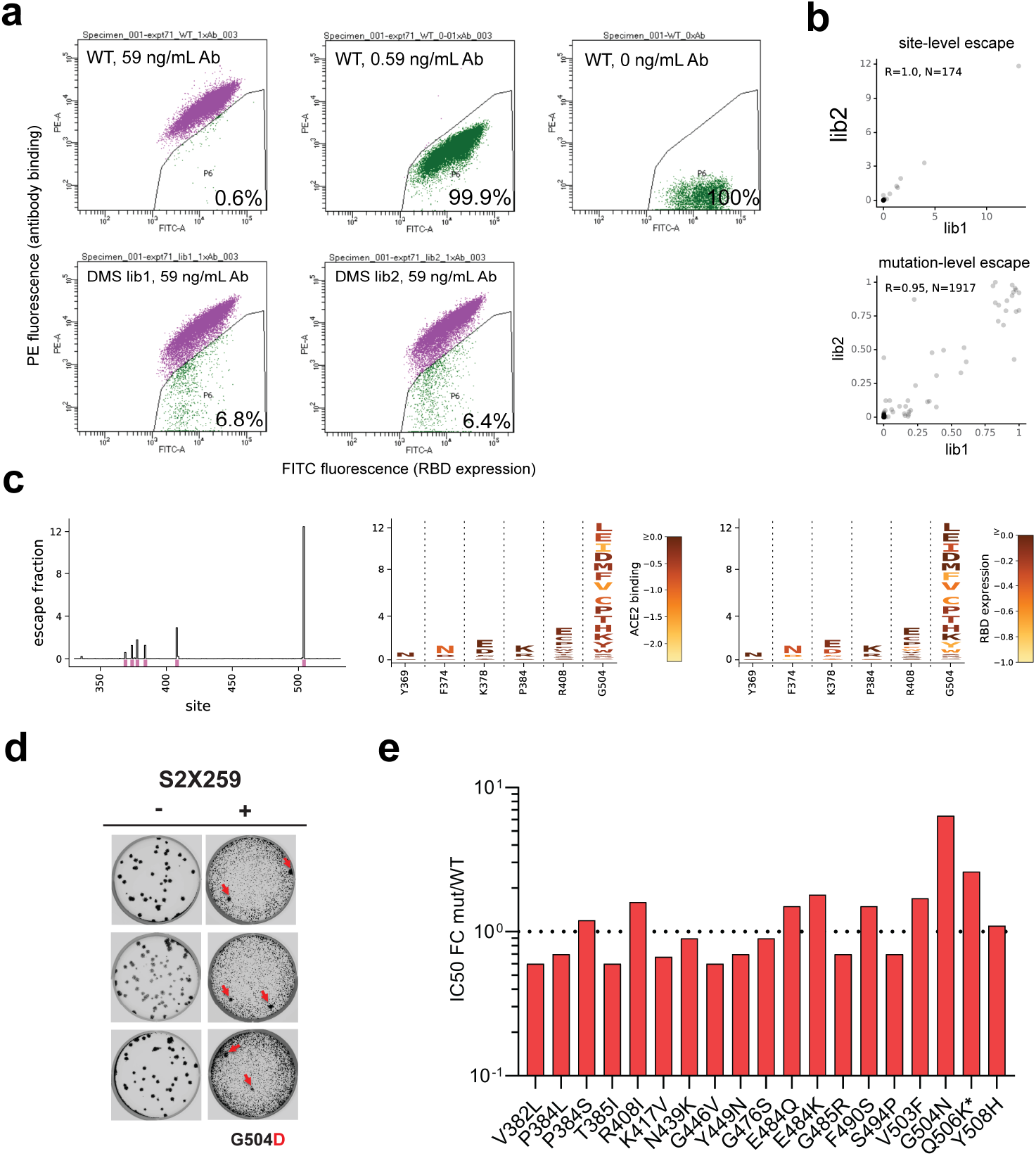
S2×259 has a high barrier for the emergence of resistance mutants. **a**, FACS gates used in DMS to select escape variants. Top row: yeast controls expressing unmutated SARS-CoV-2 RBD labelled at relevant S2×259 concentrations for setting of selection gates. Bottom row: fraction of cells in SARS-CoV-2 mutant libraries falling into the antibody-escape bin. **b**, Correlation in site-level (top, sum of escape fractions for mutations at a site) and mutation-level (bottom) escape between independently generated and assayed RBD mutant libraries. **c**, Line plot of escape mutants along all positions in the SARS-CoV-2 RBD (left). Pink lines indicate sites that escape S2×259 binding illustrated at the mutation-level in logoplots (right). In logoplots, the height of a letter scales with its escape fraction. Letters are coloured according to their deleterious consequences for ACE2 binding (middle) or RBD expression (right) as determined in prior deep mutational scanning experiments^33^. **d**, Plaque assay on Vero cells with no antibody (left) or S2×259 (right) in the overlay to isolate escape mutants (red arrow). Data are representative of three independent experiments. **e**, S2×259 in vitro neutralizing activity against SARS-CoV-2 S VSV pseudotyped mutants. For each mutant the fold change of the IC50 geometric mean vs SARS-CoV-2 S WT is reported. *Q506K displayed a 10-fold reduction in viral entry in comparison to the other mutants. Results from two independent experiments are reported.

**Extended Data Fig. 7.**
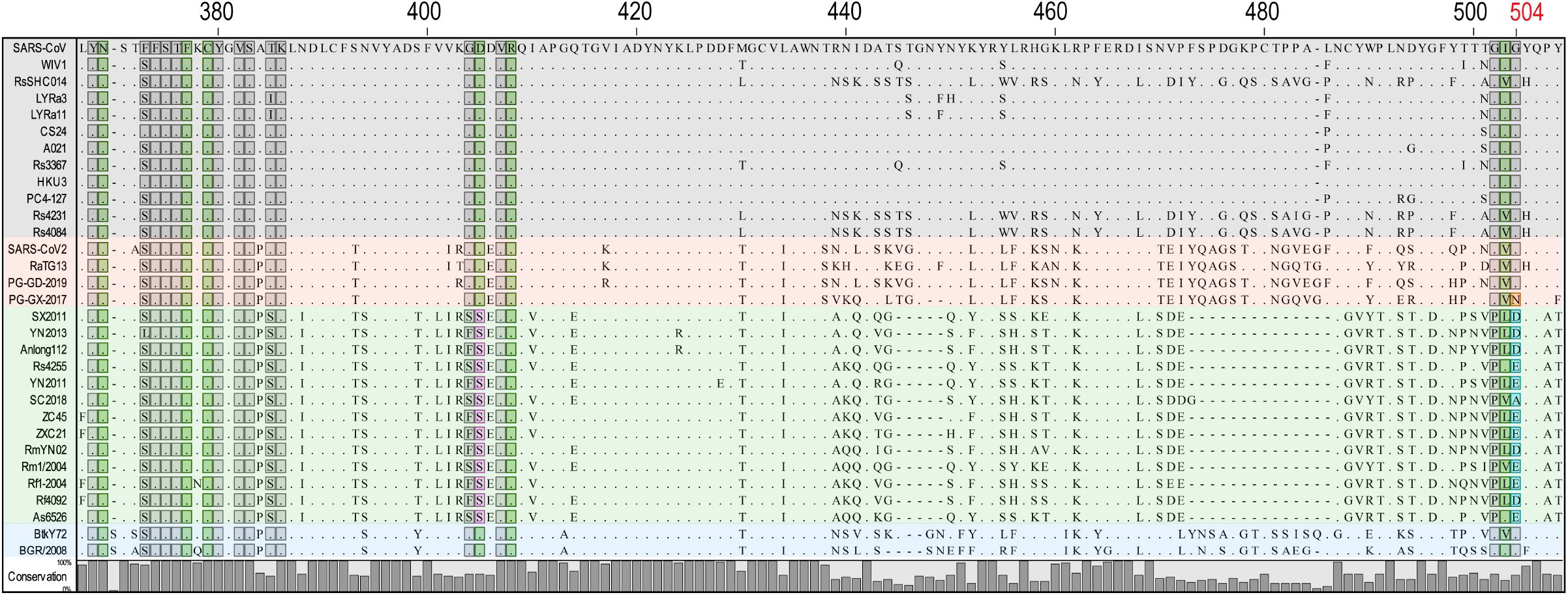
S2×259 epitope conservation across sarbecovirus clades. Protein sequence alignment of representative sarbecovirus RBDs with strictly conserved residues shown as dots. Overall conservation is represented as a bar plot. Positions are based on SARS-CoV-2 RBD. Residues determined to be important for S2×259 binding are denoted in dark green. Substitutions at positions D405 and G504 are indicated in pink and blue/orange, respectively. Additional residues representing extended epitope, are denotated grey. Different clades within the sarbecovirus subgenus are overlayed in grey (clade 1a), red (clade 1b), green (clade 2), and light blue (clade 3).

**Extended Data Fig. 8.**
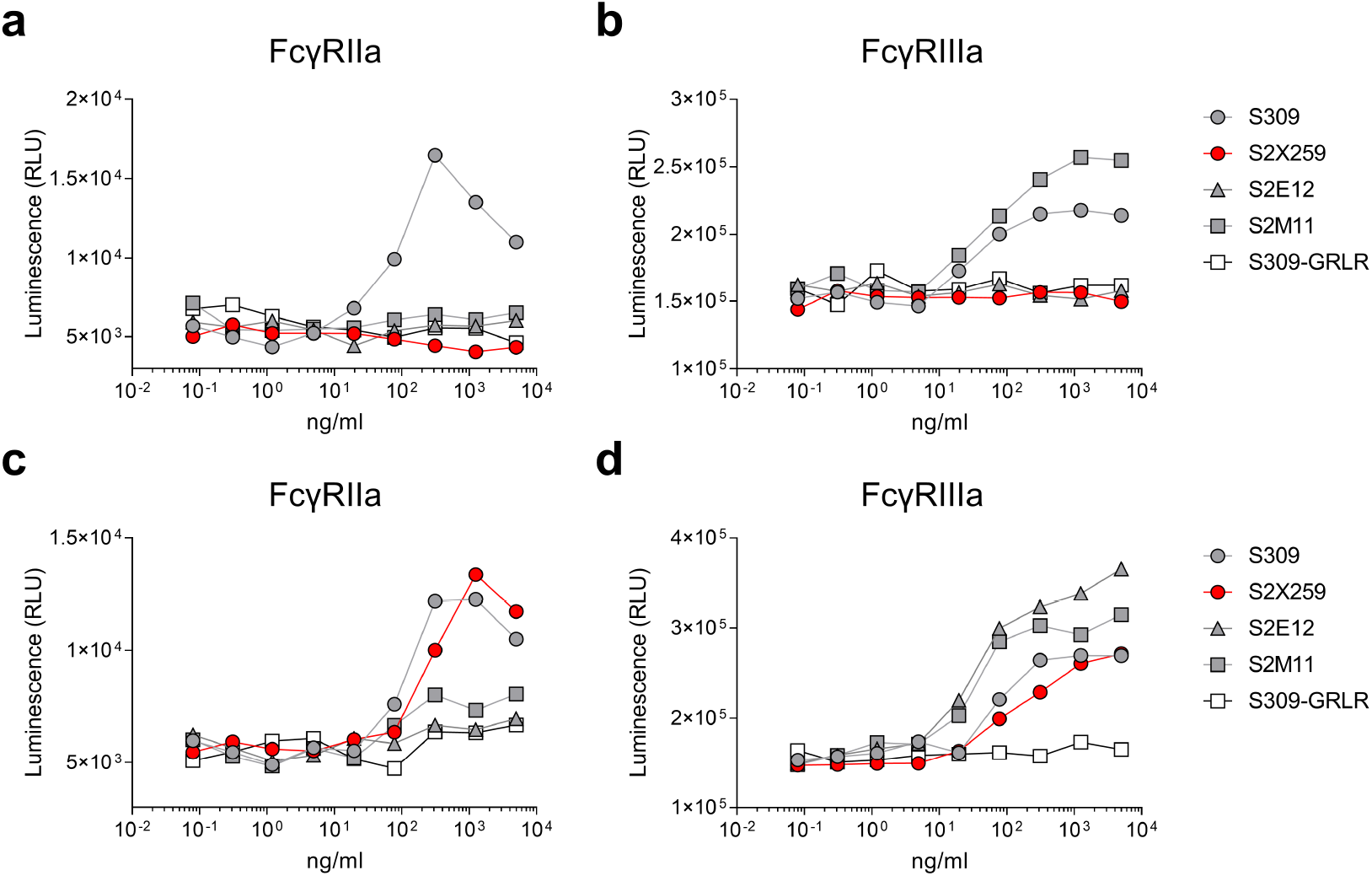
S2×259 Fc-mediated activation of FcγRIIa and FcγRIIIa *in vitro*. **a-b**, NFAT-driven luciferase signal induced in Jurkat cells stably expressing FcγRIIa H131 (a) variant or FcγRIIIa V158 (b) variant by S2×259 binding to full-length wild-type SARS-CoV-2 S on ExpiCHO target cells. **c-d**, NFAT-driven luciferase signal induced in Jurkat cells stably expressing FcγRIIa H131 (c) or FcγRIIIa V158 (d) variants by S2×259 binding to uncleavable full-length pre-fusion stabilized SARS-CoV-2 S (unable to release the S_1_ subunit) transiently expressed in ExpiCHO cells. SE12, S2M11, S309, S309-GRLR mAbs are included as controls.

**Extended Data Fig. 9.**
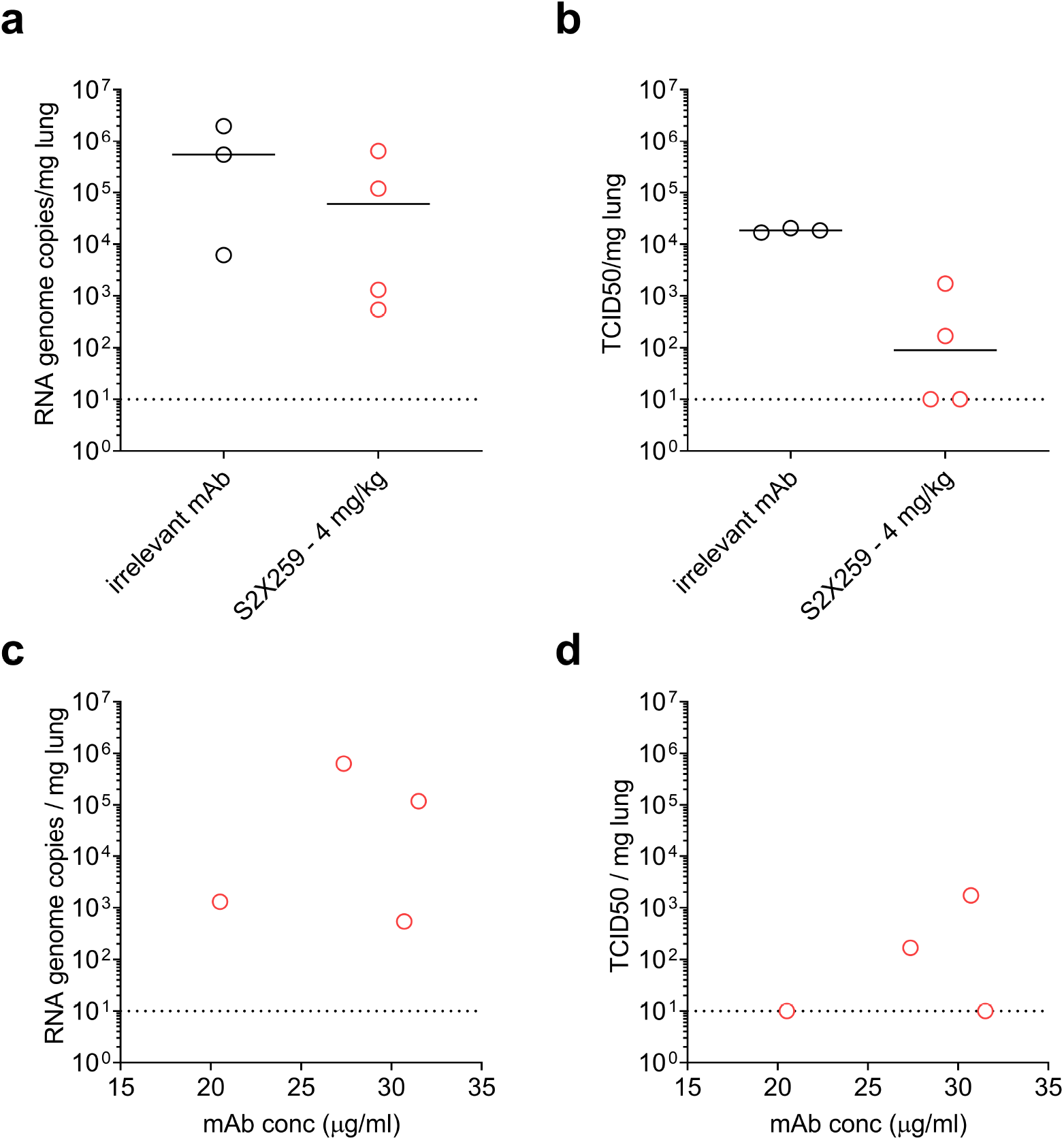
**Prophylactic administration of S2×259 mAb protects hamsters against prototypic (Wuhan-1 related) SARS-CoV-2 challenge.** **a-b**, Viral RNA loads (a) and replicating virus titres (b) in the lungs of Syrian hamsters 4 days post-intranasal infection with prototypic SARS-CoV-2. Results for one independent experiment are shown. **c-d**, Viral RNA loads and replicating virus titers in the lung 4 days post infection plotted as a function of serum mAb concentrations before infection (day 0). Irrelevant mAb n=3; S2×259 4 mg/kg n=4.

## SUPPLEMENTARY TABLES

**Table S1.** **Characteristics of S2×259. VH and VL % identity refers to V gene identity compared to germline (IMGT).**

|  |  |
| --- | --- |
| **Donor Gender** | Male |
| **Donor Age** | 52 |
| **Donor Status** | Symptomatic |
| **Days after symptoms onset** | 75 |
| **VH gene usage** | VH1-69 |
| **VH % identity** | 94.1 |
| **HCDR3** | ARGFNGNYYGWGDDDAFDI |
| **VL gene usage** | VL1-40 |
| **VL % identity** | 98.26 |
| **LCDR3** | QSYDSSLSGPNWV |

**Table S2.** CryoEM data collection and refinement statistics.

|  | SARS-CoV-2<br>S/S2X259 | SARS-CoV-2 S/S2X259<br>(local refinement) |
| --- | --- | --- |
| **Data collection and processing** |  |  |
| Magnification | 130,000 | 130,000 |
| Voltage | 300 | 300 |
| Electron exposure (e<sup>-</sup>/Å<sup>2</sup>) |  |  |
| Defocus range (μm) | 0.8-2.0 | 0.8-2.0 |
| Pixel size (Å) | 0.525 | 0.525 |
| Symmetry imposed | C3 | C1 |
| Final particle images (no.) |  |  |
| Map resolution (Å) | 3.2 | 3.5 |
| FSC threshold | 0.143 | 0.143 |
| **Validation** |  |  |
| MolProbity score | 1.08 | 0.74 |
| Clash score | 1.7 | 0.16 |
| Poor rotamers (%) | 0.38 | 5.29 |
| Ramachandran plot |  |  |
| Favored (%) | 97.1 | 97.01 |
| Allowed (%) | 2.9 | 2.74 |
| Disallowed (%) | - | 0.25 |

**Table S3.** X-ray data collection and refinement statistics.

|  |  |
| --- | --- |
|  | RBD/S2X259/S2H97<br>PDB ID: 7M7W |
| **Data collection** |  |
| Space group | P2<sub>1</sub> |
| Cell dimensions |  |
| *a*, *b*, *c* (Å) | 86.19, 66.40, 237.66 |
| $\alpha$, $\beta$, $\gamma$ (°) | 90.00, 94.34, 90.00 |
| Resolution (Å) | 63.94-2.65 (2.70-2.65) |
| *R*<sub>merge</sub> | 0.149 (2.494) |
| *I* / $\sigma I$ | 10.9 (0.8) |
| Completeness (%) | 98.6 (98.3) |
| Redundancy | 6.9 (6.8) |
| **Refinement** |  |
| Resolution (Å) | 2.65 |
| No. reflections | 73,189 |
| *R*<sub>work</sub> / *R*<sub>free</sub> | 0.221/0.271 |
| No. atoms |  |
| Protein | 16,162 |
| Ligand/ion | 28 |
| Water | 95 |
| *B*-factors |  |
| Protein | 75.86 |
| Ligand | 84.00 |
| Water | 50.09 |
| R.m.s. deviations |  |
| Bond lengths (Å) | 0.002 |
| Bond angles (°) | 0.817 |

Values for the highest-resolution shell are shown in parentheses.

**Table S4.** **Summary of nucleotide and amino acid mutations found in 18 neutralization-resistant VSV-SARS-CoV-2-S chimera plaques.**

| **Plaque** | **Nucleotide mutant** | **Amino acid Mutant** |
| --- | --- | --- |
| 1 | G1511A | G504**D** |
| 2 | G1511A | G504**D** |
| 3 | G1511A | G504**D** |
| 4 | G1511A | G504**D** |
| 5 | G1511A | G504**D** |
| 6 | G1511A | G504**D** |
| 7 | G1511A | G504**D** |
| 8 | G1511A | G504**D** |
| 9 | G1511A | G504**D** |
| 10 | G1511A | G504**D** |
| 11 | G1511A | G504**D** |
| 12 | G1511A | G504**D** |
| 13 | G1511A | G504**D** |
| 14 | G1511A | G504**D** |
| 15 | G1511A | G504**D** |
| 16 | G1511A | G504**D** |
| 17 | G1511A | G504**D** |
| 18 | G1511A | G504**D** |

